# Cooperativity between H3.3K27M and PDGFRA poses multiple therapeutic vulnerabilities in human iPSC-derived diffuse midline glioma avatars

**DOI:** 10.1101/2023.02.24.528982

**Authors:** Kasey R. Skinner, Tomoyuki Koga, Shunichiro Miki, Robert F. Gruener, Florina-Nicoleta Grigore, Emma H. Torii, Davis M. Seelig, Yuta Suzuki, Daisuke Kawauchi, Benjamin Lin, Denise M. Malicki, Clark C. Chen, Etty N. Benveniste, Rakesh P. Patel, Braden C. McFarland, R. Stephanie Huang, Chris Jones, Alan Mackay, C. Ryan Miller, Frank B. Furnari

## Abstract

Diffuse midline glioma (DMG) is a leading cause of brain tumor death in children. In addition to hallmark H3.3K27M mutations, significant subsets also harbor alterations of other genes, such as *TP53* and *PDGFRA*. Despite the prevalence of H3.3K27M, the results of clinical trials in DMG have been mixed, possibly due to the lack of models recapitulating its genetic heterogeneity. To address this gap, we developed human iPSC-derived tumor models harboring TP53^R248Q^ with or without heterozygous H3.3K27M and/or PDGFRA^D842V^ overexpression. The combination of H3.3K27M and PDGFRA^D842V^ resulted in more proliferative tumors when gene-edited neural progenitor (NP) cells were implanted into mouse brains compared to NP with either mutation alone. Transcriptomic comparison of tumors and their NP cells of origin identified conserved JAK/STAT pathway activation across genotypes as characteristic of malignant transformation. Conversely, integrated genome-wide epigenomic and transcriptomic analyses, as well as rational pharmacologic inhibition, revealed targetable vulnerabilities unique to the TP53^R248Q^; H3.3K27M; PDGFRA^D842V^ tumors and related to their aggressive growth phenotype. These include *AREG*-mediated cell cycle control, altered metabolism, and vulnerability to combination ONC201/trametinib treatment. Taken together, these data suggest that cooperation between H3.3K27M and PDGFRA influences tumor biology, underscoring the need for better molecular stratification in DMG clinical trials.

## Introduction

Diffuse midline glioma (DMG), H3 K27-altered, is a relatively new diagnostic entity defined in the 2021 World Health Organization Classification of Tumors of the Central Nervous System (Louis et al. 2021). This diagnosis occurs primarily in children and includes tumors previously referred to as “diffuse intrinsic pontine glioma”, or “diffuse midline glioma, H3 K27M-mutant.” Tumors in this category are characterized by K27M mutations in genes encoding variants of histone H3: *H3F3A* (H3.3K27M) or, less commonly, *HIST1H3B* (H3.1K27M) (Louis et al. 2016). H3.3K27M in particular defines this class of DMG because it occurs in approximately 60% of brainstem and non-brainstem midline high-grade gliomas in children (Mackay et al. 2017) and correlates strongly with poor patient outcomes (Sturm et al. 2012). DMG are poor candidates for surgery and radiation due to their diffusely invasive nature and sensitive location, resulting in a dismal median survival of 11 months (Hoffman et al. 2018).

H3.3K27M is associated with regional decreases in H3K27 trimethylation (H3K27me3) (Chan et al. 2013; Lewis et al. 2013), a histone posttranslational modification associated with transcriptional repression (Cao et al. 2002). Physiologic H3K27me3-mediated transcriptional control is vital for the appropriate regulation of many processes in central nervous system development, including maintenance of stem cell pluripotency and glial cell fate specification and proliferation (Boyer et al. 2006; Lee et al. 2006). Therefore, the epigenetic rewiring induced by H3.3K27M dysregulates these sensitive processes, promoting gliomagenesis (Berdasco et al. 2010). Despite the progress made in elucidating the impact of H3.3K27M on cellular biology, the mechanisms by which it leads to DMG remains to be found. Further, drugs that show preclinical promise have failed to extend median overall survival in clinical trials. As a result, H3K27-altered DMG remains a leading cause of brain tumor-related death in children (Ostrom et al. 2020).

One potential reason for trial failures is the need for preclinical models that accurately recapitulate DMG genetics. H3.3K27M almost always co-occurs with alterations in tumor suppressors and/or oncogenes, including *TP53, PDGFRA, PIK3CA*, and *NF1* (Mackay et al. 2017). Mouse models have shown that H3.3K27M expression alone is not sufficient to induce gliomas and that adding PDGFRA overexpression or TP53 loss potentiates tumorigenesis (Cordero et al. 2017; Funato et al. 2014; Misuraca et al. 2016), suggesting functional cooperation of these alterations. While available models of H3.3K27M DMG utilize *PDGFRA* or *TP53* mutations, little is known about how these alterations interact with H3.3K27M to affect processes in both normal and malignant cells. Thus, genetically faithful models of DMG with H3.3K27M and other cooccurring mutations are needed to predict the success of therapies in patients more accurately.

Genetically engineered human induced pluripotent stem cells (iPSC) have proven to be an effective, highly customizable system to model gliomagenesis in a species-specific manner, allowing for longitudinal analysis of human tumor biology from potential cell(s) of origin (Koga et al. 2020; Miki et al. 2022; Parisian et al. 2020). Here, we employ an iPSC-based system to show that *TP53* loss- and *PDGFRA* gain-of-function mutations, respectively, cooperate with heterozygous, chromosomal H3.3K27M to potentiate DMG tumor growth and alter cellular metabolism and intracellular signaling. Further, we demonstrate the utility of these models in preclinical drug testing.

## Results

### Neural progenitors harboring TP53 and H3.3K27M mutations form tumors resembling DMG

To generate a genetically faithful model of H3.3K27M DMG, we introduced mutations into *TP53* and *H3F3A* in human iPSC by CRISPR/Cas9 genome editing (Fig. 1A). *TP53^R248Q^* is a dominant negative mutation in the DNA binding domain of the TP53 protein, mimicking the loss of TP53 function commonly seen in H3.3K27M tumors (Hanel et al. 2013). Since patient tumors are H3F3A^K27M^ heterozygous, iPSC clones harboring heterozygous H3F3A^K27M^ mutations were selected to better recapitulate the genetics of the disease than models relying on H3.3K27M overexpression (Schwartzentruber et al. 2012). Because neural progenitor cells (NPC) are a likely cell of origin for gliomas, including DMG, we chemically differentiated TP53^R248Q^ (henceforth referred to as “T”) and TP53^R248Q^; H3.3K27M (henceforth referred to as “TH”) iPSC into induced NPC (iNPC) (Reinhardt et al. 2013). RT-qPCR showed decreased expression of iPSC markers (*NANOG, OCT4*) and increased expression of the NPC marker *PAX6* in iNPC relative to iPSC (Fig. 1B). We verified *TP53* and *H3F3A* genotypes of resultant T and TH iNPC via Sanger sequencing (Fig. 1C).

**Figure 1.**
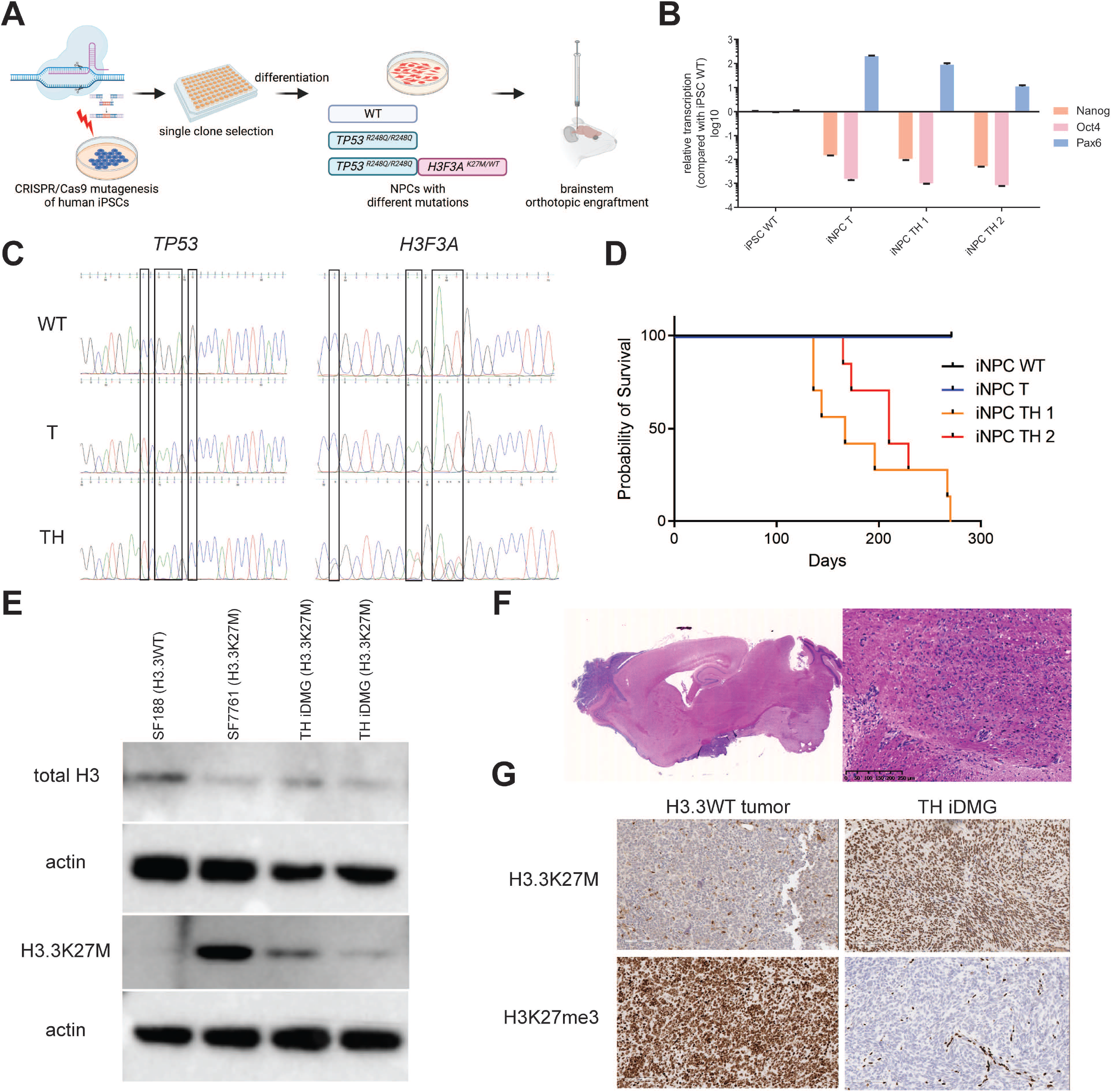
iPSC-derived neural progenitors harboring TP53^R248Q^ and H3.3K27M generate high-grade gliomas with H3.3K27M hallmarks *in vivo*. (**A**) Schematic of generation of iDMG harboring TP53^R428Q^ (T), with and without H3.3K27M (H). (**B**) qRT-PCR of iPSC (*NANOG* and *OCT4*) and neural progenitors (*PAX6*) marker RNA expression in iPSC and iNPC. (**C**) *TP53* and *H3F3A* Sanger sequencing showing the location of alterations in wild-type, T, and TH iNPC. (**D**) Kaplan-Meier curves showing survival of immunodeficient mice orthotopically engrafted with wild-type, T, or TH iNPC. n=4 animals for WT iNPC, n=4 animals for T iNPC, n=7 animals each for two TH iNPC clones. Median survival is not significantly different between TH iNPC clones (log-rank test, p=0.48). (**E**) Western blots of total H3 and H3.3K27M expression in established pediatric glioma cell lines and two TH iDMG tumors. (**F**) 0.4x (left) and 10x (right, scale bar = 250 μm) views of a H&E-stained sections from a TH iDMG tumor. (**G**) Immunohistochemical staining for H3.3K27M (top) and H3K27me3 (bottom) of tumors from mice engrafted with GBM39 (left, H3.3WT) and TH iNPC (right). Scale bars = 100 μm.

We next investigated the tumorigenic potential of both T and TH iNPC by engrafting them into the brains of immunocompromised mice. TH iNPC formed orthotopic brain tumors with median survival of 167 days for one clone and 210 days for the second. Neither wild type nor T iNPC formed tumors within this experimental time frame (Fig. 1D). TH tumors retain expression of H3.3K27M via Western blot, with established cell lines derived from H3.3WT (SF188) and H3.3K27M (SF7761) pediatric gliomas used as negative and positive controls. TH tumors from two iNPC clones expressed mutant H3.3 at similar levels relative to total H3 (Fig. 1E). Hematoxylin & eosin (H&E) staining of TH tumors showed diffusely invasive masses in the brainstem or cerebellum. These masses harbored mitotically active cells with nuclear atypia, in keeping with histologic features of high-grade gliomas (Fig. 1F). Immunohistochemistry confirmed almost universal positivity of neoplastic cells for H3.3K27M; a control tumor generated from an adult, H3.3 wild-type glioblastoma patient-derived xenograft (PDX, GBM39) was H3.3K27M-negative (Fig. 1G, top) (Vaubel et al. 2020). TH tumors also showed markedly decreased H3K27me3 staining in comparison with GBM39, consistent with the decrease in H3K27me3 deposition characteristic of H3.3K27M DMG (Fig. 1G, bottom). Based on these findings, we concluded that TH tumors generated from iPSC-derived NPC resemble H3.3K27M DMG.

### PDGFRA activation enhances tumorigenesis

We generated TH tumors to study the transcriptomic and epigenomic effects of H3.3K27M. However, the inability of T iNPC to form tumors in mice limited our ability to isolate the effect of H3.3K27M on the tumor transcriptome. To better recapitulate the genetic heterogeneity observed in DMG and to generate isogenic H3.3WT tumors for comparison to TH tumors, we overexpressed a constitutively activating mutation found in DMG (Paugh et al. 2013), PDGFRA^D842V^, in both T (TP) and TH (THP) iNPC (Fig. 2A). PDGFRA^D842V^ enhanced the transcriptomic differences between H3.3K27M and H3.3WT iNPC. Unsupervised hierarchical clustering grouped TP53^R248Q^;PDGFRA^D842V^ by H3.3 status in contrast to TP53^R248Q^ iNPC (Fig. 2B). The 135 differentially expressed genes (DEG) (Q<0.05, Supplemental Tables) between THP and TP iNPCs had significant overlap with gene sets related to cancer in general and glioma in particular (Fig. S1A&B). These genes also were enriched in pathways related to both early development and growth- and invasion-related processes (Fig. S1C), leading us to hypothesize that PDGFRA^D842V^ cooperates with H3.3K27M to create an optimal cellular environment for malignant NPC transformation.

**Figure 2.**
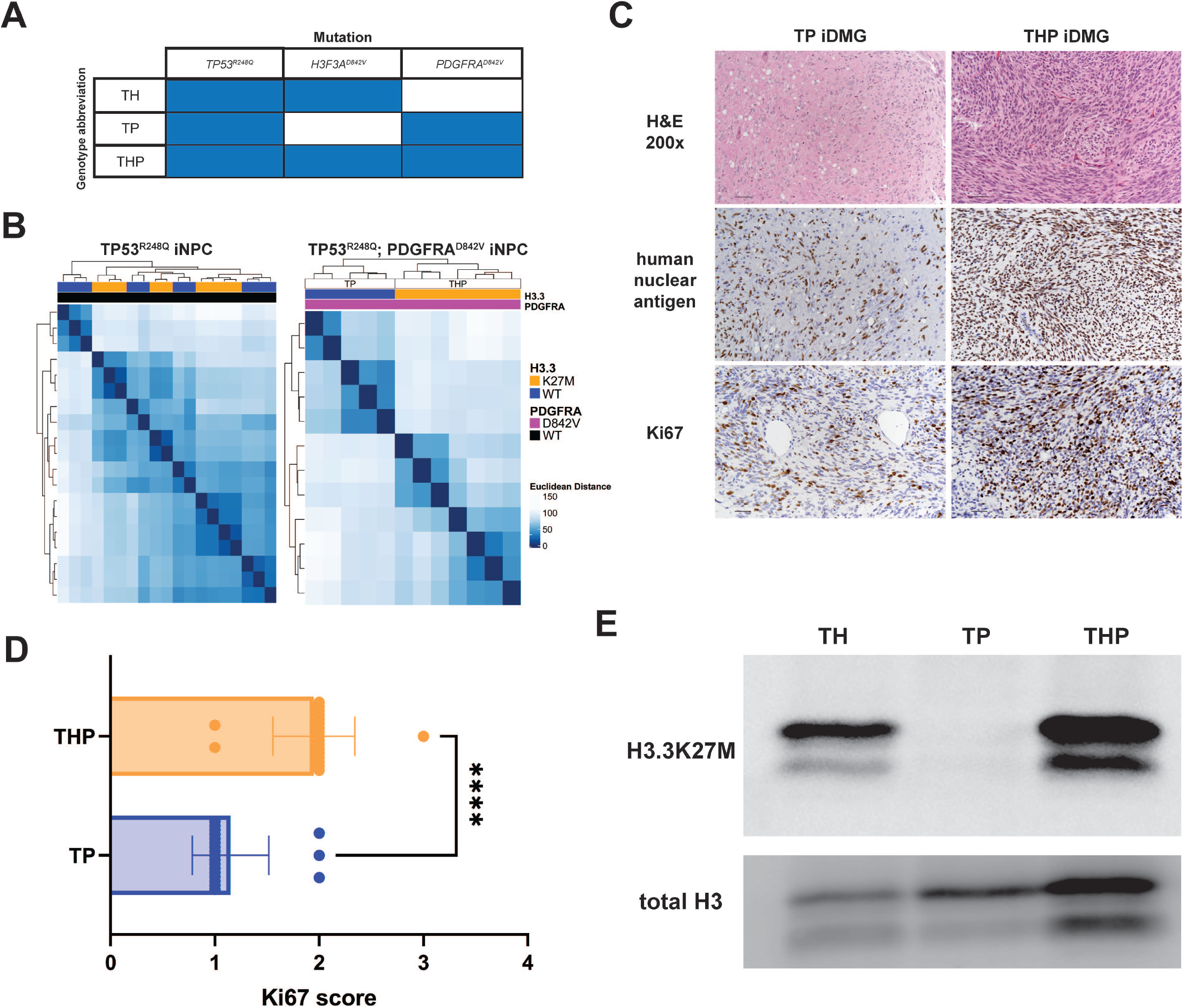
PDGFRA^D842V^ enhances transcriptome differences and potentiates tumorigenesis in H3.3K27M iDMG. (**A**) iDMG genotypes and abbreviations. T=TP53^R248Q^; H=H3F3AK27M; P=PDGFRA^D842V^. (**B**) Unsupervised hierarchical clustering based on Euclidean distance between RNA-seq samples of iNPC tumor cells with (right) and without (left) PDGFRA^D842V^ shows enhanced clustering by H3.3 status. (**C**) From top down: H&E (scale bar = 50 μm), anti-human nuclear antigen (scale bar = 50 uM), and anti-Ki67 (scale bar = 50 μm) stained sections from representative THP and TP iDMG. (**D**) Semi-quantitative scoring of Ki67-positive neoplastic cells. Two tumors per genotype were scored. t-test, ****=p<0.0001. (**E**) Western blots of H3.3K27M and total H3 in whole cell lysates from dissociated iDMG cultured on Cultrex.

To determine the impact of this cooperation on gliomagenesis, we implanted TH, TP, and THP iNPCs into the brainstems of mice. THP (median=27 days) or TP (median=27 days) tumor bearing mice survived significantly shorter (log-rank test, p<0.0001) than those with TH tumors (median=203 days) (Fig. S2A). H&E-stained sections of TP and THP tumors showed poorly demarcated, densely hypercellular, diffusely infiltrative masses in the brainstem, midbrain, and/or cerebellum. High-grade glioma hallmarks such as mild to moderate nuclear pleomorphism, anisokaryosis, and occasional to frequent mitoses were evident in THP tumors. TP tumors were less cellular but showed similar nuclear pleomorphism (Fig. 2C, top). Both showed evidence of leptomeningeal tumor spread. THP showed greater Ki67 positivity among the neoplastic cells (defined by positive staining for human nuclear antigen) than TP tumors (Fig. 2C, bottom). Semi-quantitative Ki67 analysis confirmed that THP were significantly more proliferative than TP tumors (Fig. 2D). Thus, although overall survival for the mice bearing THP and TP tumors was equivalent, histologic findings suggest that THP tumors are more aggressive and proliferative than their TP counterparts, supporting our hypothesis that H3.3K27M and PDGFRA^D842V^ cooperate to enhance tumorigenesis.

To further investigate the mechanisms of these phenotypic differences, we dissociated TH, TP, and THP tumors and cultured them on Matrigel in serum free media, where they grew robustly (Fig. S2B&C). We confirmed that these tumor-derived cultures (hereafter referred to as iDMG to reflect their resemblance to the human disease) continued to express similar amounts of H3.3K27M relative to total H3 (Fig. 2E). Thus, TH, TP, and THP iDMG have the histologic characteristics of high-grade gliomas (TP would be classified as HGG instead of DMG in patients, but all are referred to here as iDMG for convenience), making them an ideal model system to study the individual and cooperative effects of *H3F3A* and *PDGFRA* mutations on an isogenic background.

### TP53^R248Q^;H3.3K27M;PDGFRA^D842V^ (THP) iDMG recapitulate hallmarks of the human disease

We next investigated whether THP iDMG contain the hallmarks of H3.3K27M expression as delineated by previous studies in human and mouse tumor models. Because this mutation is associated with epigenetic rewiring and related changes in gene expression, we performed Cleavage Under Targets and Release Using Nuclease (CUT&RUN) and RNA sequencing (RNA-seq) on cultured THP and TP iDMG. First, we confirmed overall reduction of H3K27 trimethylation (H3K27me3), a characteristic of H3.3K27M tumors across models and species (Fig. 3A). H3K27me3 CUT&RUN was concordant with the Western blot results, showing that most loci heavily occupied by H3K27me3 in TP are lost in THP iDMG (Fig. 3B, left). This finding was supported by quantitative normalization using spike-in *E. coli* DNA (Fig. 3B, right). In addition to this overall decrease in H3K27me3 occupancy, THP iDMG capture the complexity of epigenetic changes seen in patient tumors. For example, loci such as *CDKN2A* retain H3K27me3 in THP relative to TP iDMG, which correlates with reduced *CDKN2A* RNA expression and is consistent with previous findings in patient-derived cell lines (Fig. S3A) (Harutyunyan et al. 2019; Mohammad et al. 2017). We found a concomitant reduction in H3K4 trimethylation (H3K4me3) and H3K27 acetylation (H3K27ac), both of which are associated with transcriptional activation, at the *CDKN2A* locus in THP vs. TP iDMG, supporting the idea that H3.3K27M-induced epigenetic remodeling is complex and not limited to H3K27me3 (Fig. S3A). Additionally, THP cells show reduced spreading of H3K27me3 from retained peaks, a phenomenon linked to H3.3K27M-mediated regulation of gene expression (Fig. S3B) (Harutyunyan et al. 2019).

**Figure 3.**
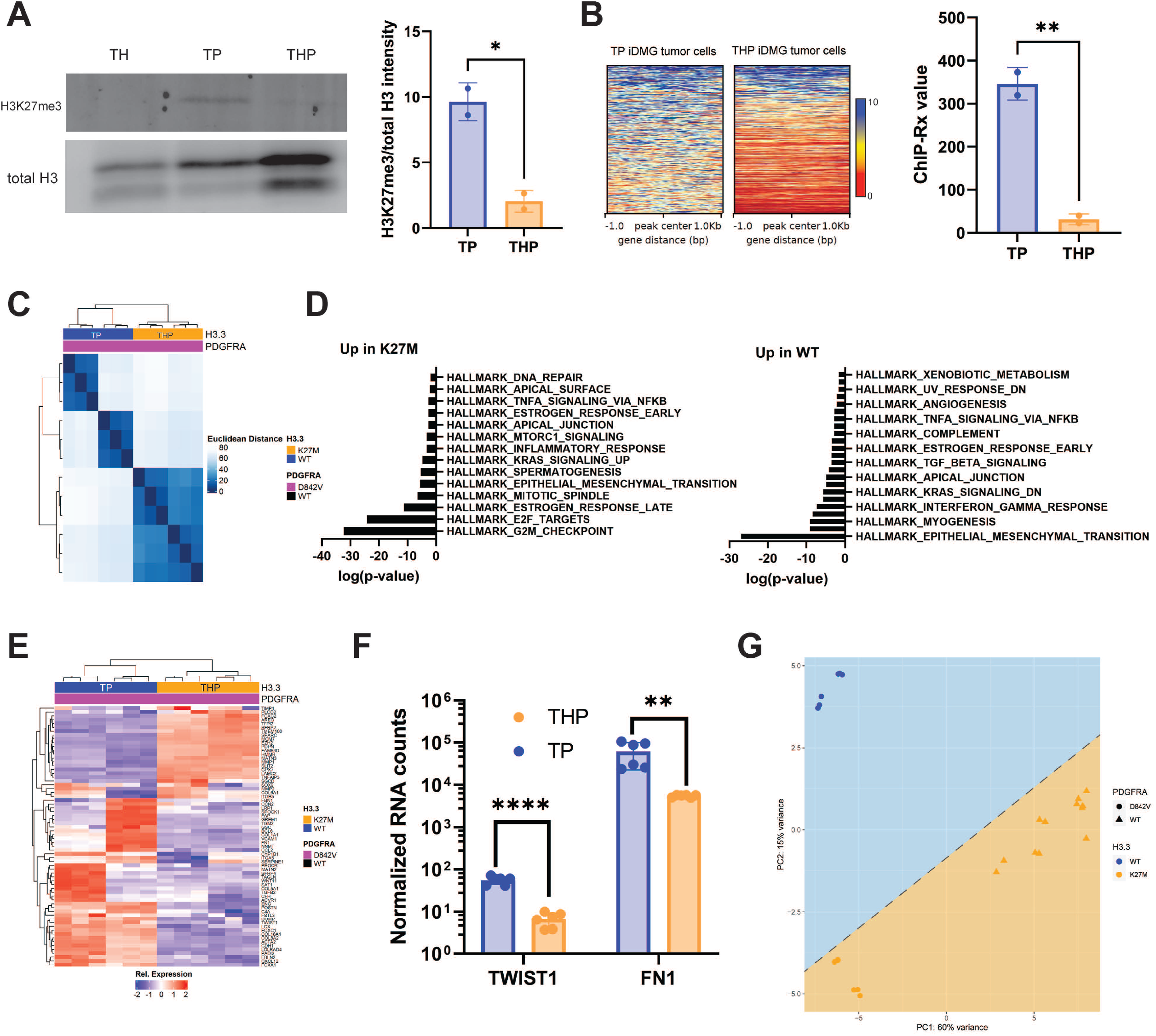
Primary tumor neurospheres generated from PDGFRA^D842V^ H3.3K27M iNPC resemble H3.3^K27M^ DMG. (**A**) Representative Western blot (left) and quantification (right, n=2 biological replicates per genotype) of H3K27me3 relative to total H3 shows a global loss of H3K27me3 in H3.3K27M tumor cells relative to wild type. t-test, *=p<0.05. (**B**) (Left) H3K27me3 occupancy in TP and THP iDMG cells. The top H3K27me3-occupied regions in TP cells are generally lost in THP cells. (Right) ChIP-Rx quantification of total H3K27me3 using E. coli spikein DNA read counts for TP and THP cells (see methods for detailed information). t-test, **=p<0.01. (**C**) Unsupervised hierarchical clustering of TP and THP iDMG tumor cells based on Euclidean distance between RNA-seq samples. (**D**) Statistical significance of overlaps (hypergeometric test) between genes significantly up- and down-regulated in THP vs TP iDMG tumor cells in MSigDB Hallmark gene sets. (**E**) Unsupervised hierarchical clustering of THP and TP iDMG tumor cells based on normalized expression of epithelial-to-mesenchymal transition genes. (**F**) Normalized *TWIST1* and *FN1* RNA read counts in THP vs. TP iDMG tumor cells (Welch’s t-test, ****=q<0.0005, **=q<0.05). (**G**) PCA plot of iDMG clustered on DEG between H3.3K27M and non-K27M pHGG patient-derived tumor and cell line samples.

THP transcriptomes are also distinct from TP iDMG; unsupervised hierarchical clustering showed tight grouping by H3.3 status (Fig. 3C). To interrogate the genes and pathways responsible for this difference, we performed differential expression analysis between TP and THP iDMG and calculated the overlap of the 539 DEG (Q<0.05, Supplemental Tables) with MSigDB Hallmark gene sets. THP cells upregulate genes associated with cell cycle progression and intracellular signaling, consistent with the aggressive, proliferative nature of these tumors. We also found significant overlap between both up- and down-regulated DEG and the Hallmark epithelial to mesenchymal transition (EMT) gene set (Fig. 3D). Differential expression of EMT-related genes has been shown to be a feature of H3.3K27M DMG. Genes typically expressed post-EMT are downregulated in H3.3K27M tumors (Sanders et al. 2020). We clustered iDMG on a list of EMT-related genes (Sanders et al. 2020) and showed that H3.3 status highly influences their expression (Fig. 3E). THP relative to TP iDMG show decreased expression of *TWIST1* and *FN1*, genes that are expressed post-EMT in normal tissue and downregulated in H3.3K27M tumors (Fig. 3F) (Sanders et al. 2020). This finding contrasts with increased *TWIST1* and *FN1* expression in adult GBM compared to normal tissue, highlighting the unique biology of H3.3K27M DMG (Brennan et al. 2013; Elias et al. 2005).

Finally, to directly compare iDMG to patient tumors, we obtained previously published RNA-seq, whole exome sequencing, and whole genome sequencing from a patient-derived panel of 62 pediatric high-grade glioma (pHGG) cell lines and 138 pHGG tumors (Carvalho et al. 2020; Izquierdo et al. 2022; Mackay et al. 2017; Mackay et al. 2018) (Fig. S4A). We identified 133 DEG (Q<0.05, Supplemental Tables) between H3.3K27M and non-H3.3K27M samples in this patient-derived panel. This gene list clustered iDMG by both H3.3 and PDGFRA status on principal component analysis (PCA) (Fig. 3G). We obtained similar clustering using H3.3K27M vs. non-H3.3K27M DEG in separate patient-derived cell line (Fig. S5A) and tumor (Fig. S6A) cohorts. While combined PCA of the patient-derived panel and iDMG did not show a clear separation on *H3F3A* status, this was also the case with the patient samples alone and likely reflects the genetic complexity of these samples (Fig. S4B). When we compared DEG between H3.3K27M and non-H3.3K27M in iDMG and the patient-derived panel, we found a significant overlap of 21 DEG (p<6.58e-10, hypergeometric test) between the two groups (Fig. S4C). Overlapping genes were significantly associated with early embryonic development and loci normally occupied by H3K27me3 in embryonic stem cells (Fig. S4D). We performed the same analysis on the patient-derived cell lines (Fig. S5B&C) and patient tumors (Fig. S6B&C) separately and found that they also shared with iDMG H3.3 genotype-dependent DEG that are significantly associated with PRC2 and EED targets. This finding suggests that in both iDMG and patient samples, H3.3K27M is associated with differential expression of genes marked by H3K27me3 either during development or in mature cells. Thus, the transcriptomic features that THP iDMG share with H3.3K27M patient samples are highly specific to this histone mutation. Taken together, these data validate THP iDMG as a model of the human disease, with key molecular phenotypes observed in patient tumors.

### The transition from iNPC to tumor is marked by conserved IL6-JAK-STAT expression

Mechanistic dissection of gliomagenesis through direct comparison of tumor cells to their iNPC cells of origin is a major advantage of iDMG. Having confirmed that THP iDMG recapitulates well the transcriptomic and epigenetic landscapes of H3.3K27M DMG, we next sought gene expression programs that define tumorigenesis driven by PDGFRA, H3.3K27M, or both. To do so, we compared RNA-seq data from TH, TP, and THP iDMG and the iNPC from which these tumors originated. Surprisingly, all three genotypes shared 1400 DEG between iNPC and iDMG (Q<0.05, Supplemental Tables), indicating that some pathways involved in malignant transformation are conserved regardless of driver mutation profile (Fig. 4A). Signaling programs including TNFA and IL6-JAK-STAT3 are among the most significantly enriched gene sets for tumorigenesis in all three studied genotypes using gene set over-representation analysis on 1400 common tumorigenesis genes (Fig. 4B). Because JAK/STAT has been linked to EMT in DMG and other cancers, we further investigated its role in tumorigenesis using iDMG (Olaciregui et al. 2018; Wang et al. 2018; Zhong et al. 2020). Unsupervised hierarchical clustering on members of the Hallmark IL6-JAK-STAT3 gene set showed that most pathway genes are upregulated in iDMG compared to iNPC (Fig. 4C). Some genes show higher expression in one genotype relative to the others, but important markers of JAK/STAT pathway activation including *IL6, OSMR, STAT3*, and *SOCS3* are consistently upregulated in tumor cells across genotypes (Fig. 4D).

**Figure 4.**
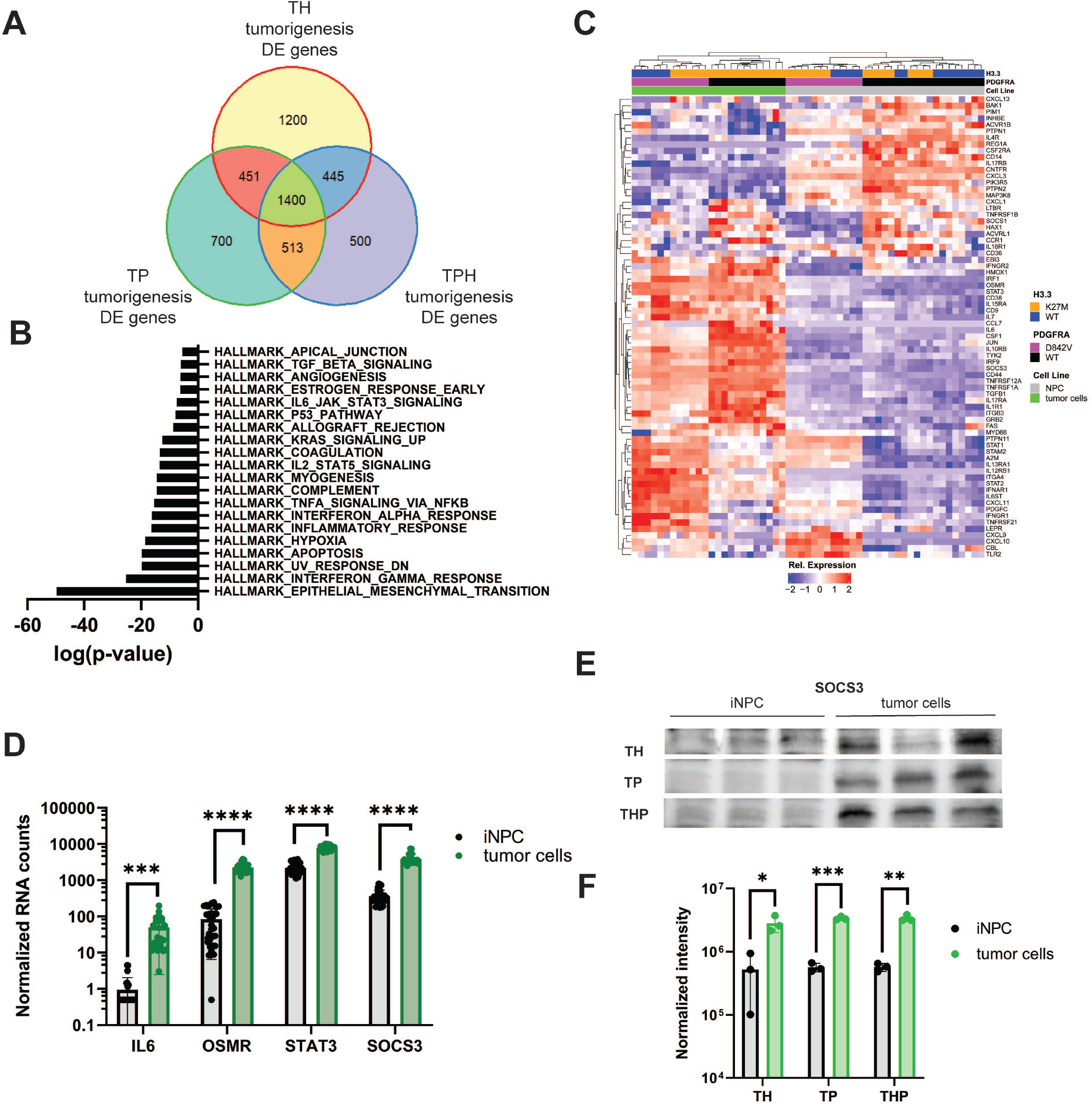
IL6-JAK-STAT3 pathway genes are associated with iDMG tumorigenesis regardless of PDGFRA and H3.3 status. (**A**) Venn diagram of DEG (Q<0.05) between iNPC cells and iDMG tumor cells for TH, TP, and THP genotypes. (**B**) Statistical overlap of the 1400 common DEG from the center of the Venn diagram above with MSigDB HALLMARK gene sets. (**C**) Unsupervised hierarchical clustering of TH, TP, and THP NPC and iDMG on Hallmark IL6-JAK-STAT3 pathway genes. (**D**) DESeq2 normalized RNA counts of key IL6-JAK-STAT3 pathway genes (Welch’s t-test, ****=p<0.00001, ***=p<0.00005). (**E**) Western blots of SOCS3 expression THP NPC and iDMG (3 biological replicates each cell type). (**F**) Quantification of SOCS3 expression via Western blot for TH, TP, and THP iNPC vs. iDMG (n=3 biological replicates per genotype per cell type, Welch’s t-test, *=q<0.05, **=q<0.005, ***=q<0.0005). Bands were normalized to total protein per lane.

SOCS3 is a negative regulator of IL6-JAK-STAT3 signaling upregulated specifically by STAT3 activation (Yoshimura et al. 2007), suggesting that pathway members are indeed more active in iDMG than in iNPC rather than simply being more highly transcribed. We confirmed increased SOCS3 protein expression by Western blot using total lane protein normalization. Significantly higher levels of SOCS3 protein were found in iDMG tumor cells vs. iNPC across all three genotypes (Fig. 4E&F). However, STAT3 phosphorylation was only increased in TH iDMG relative to iNPC, suggesting that high SOCS3 may have suppressed STAT3 phosphorylation in other genotypes (data not shown). These results implicate IL6-JAK-STAT3 as a key conserved player in DMG tumorigenesis regardless of *H3F3A* or *PDGFRA* genotype and showcase the unique ability of iDMG models to act as a temporal window into the tumorigenic process.

### Epigenetic upregulation of amphiregulin is unique to THP iDMG and associated with proliferation

Having found a common tumorigenesis-associated pathway among genotypes, we next asked whether there are epigenetically regulated genes unique to the combination of PDGFRA^D842V^ and H3.3K27M. We identified genes upregulated in THP vs. TP iDMG with large corresponding changes in the distribution of activating and repressive epigenetic marks using Intepareto, a package that maximizes concordance among epigenetic and transcriptomic changes in RNA-seq and CUT&RUN data. Amphiregulin (*AREG*), a low affinity ligand for epidermal growth factor receptor (*EGFR*), was among the top genes and is a pertinent target due to the prevalence of *EGFR* mutations or amplification in high-grade gliomas (Fig. 5A). Loss of H3K27me3 was evident across the *AREG* gene body and gain of both H3K27 acetylation (H3K27ac) and H3K4 trimethylation (H3K4me3) was primarily evident at transcription start sites (Fig. 5B). Loss of a repressive and gain of activating histone modifications correlate with the approximately 85-fold change in *AREG* expression between THP and TP iDMG (Fig. S7A). High *AREG* RNA expression was seen only in THP iDMG, suggesting that this change is not driven by H3.3K27M alone (Fig. 5C). Significantly higher levels of AREG protein were also found in both whole cell lysate and cell culture media from THP than either TH or TP iDMG (Fig. 5D). Therefore, high expression and secretion of AREG are unique to THP iDMG.

**Figure 5.**
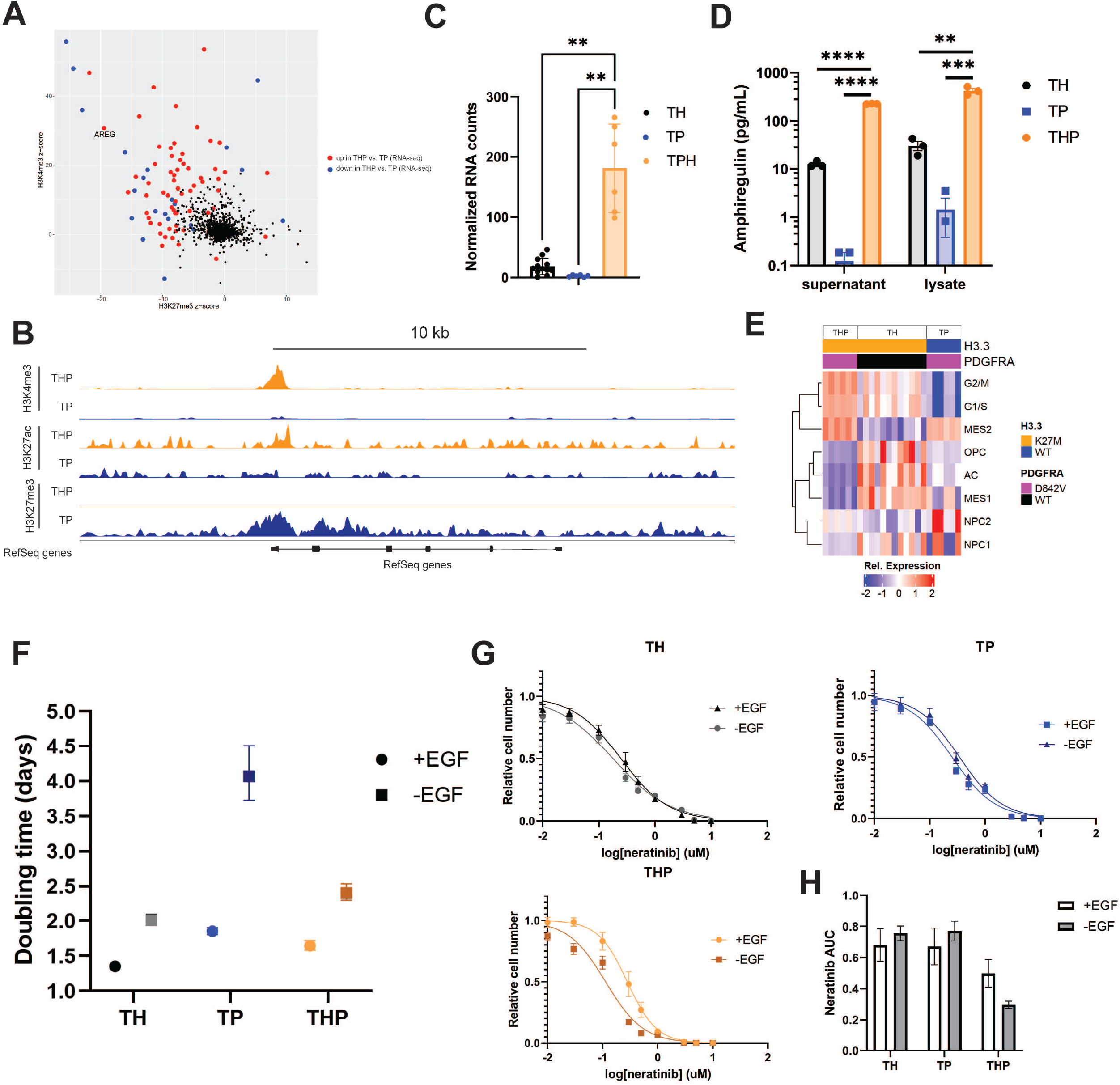
THP iDMG are uniquely reliant on AREG-mediated proliferation. (**A**) IntePareto plot showing genes with the most concordant differences in H3K27ac, H3K27me3, and H3K4me3 distribution and RNA expression between THP and TP iDMG cells. (**B**) Bigwig tracks from CUT&RUN data at the AREG locus show loss of H3K27me3 and gain of H3K4me3 and H3K27ac in THP cells relative to TP. The y-axis is the same between genotypes for each histone modification. (**C**) Normalized AREG RNA-seq counts are significantly elevated in THP vs. TH or TP iDMG cells (Welch’s t-test, p<0.005). (**D**) Amphiregulin concentration in cell culture supernatant and whole cell lysates measured via ELISA (t-test, **=q<0.01, ***=q<0.001, ****=q<0.0001). (**E**) ssGSEA heatmap of iDMG relative enrichment scores for cell type-specific gene sets. (**F**) Doubling times for TH, TP, and THP iDMG cells in the presence and absence of EGF/bFGF determined via five-day in vitro proliferation. Doubling times are significantly different between +EGF and −EGF conditions for each genotype (extra sum of squares F-test, p<0.0001). (**G**) Neratinib dose response curves for TH, TP, and THP iDMG cells in vitro. Relative cell numbers were normalized to zero-drug, 1% DMSO wells. Neratinib IC50 is significantly different between the +EGF and −EGF conditions for THP only (extra sum of squares F-test, p<0.01). n=9 for each drug dose (3 experiments with 3 technical replicates per dose each). (**H**) Area under the neratinib dose response curves in G. Error bars represent SEM.

In normal development and non-CNS neoplasms, *AREG* can act as a growth factor and stimulate proliferation, migration and invasion, and therapy resistance (Busser et al. 2011). THP iDMG have the highest scores for cycling cell gene sets, while TH and TP iDMG more closely resemble less proliferative cell types (Fig. 5E) upon ssGSEA using brain cell type marker genes. Among genes uniquely upregulated in THP vs. TH or TP iDMG were important regulators of G1/S and/or G2/M cell cycle transitions, including *CDK1* and *CDK2* (Fig. S7B). These data are consistent with the highly proliferative nature of THP tumors observed by histology (Fig. 2C&D). Although the role of *AREG* has not been specifically defined in DMG, it increases cell cycle progression via the transcription factor *FOXM1* in other cancers (Stoll et al. 2016). Therefore, its role in THP iDMG may be associated with upregulation of mitogenesis.

Since amphiregulin is an EGFR ligand, we hypothesized that differential activation of and dependence on EGFR signaling could underlie the upregulation of cell cycle genes that characterize THP iDMG. *AREG* was the most significantly overexpressed of all EGFR ligands in THP iDMG versus the other genotypes, indicating that any genotype effect on the EGFR pathway is specific to AREG-dependent signaling (Fig. S7C). Significantly higher *EGFR* RNA expression in THP vs. TP iDMG also suggests that manipulation of EGFR signaling might have a disproportionate impact on this genotype (Fig. S7D). Growth inhibition experiments revealed no difference in sensitivity to the irreversible EGFR tyrosine kinase inhibitor neratinib between TH, TP, and THP iDMG *in vitro* (Fig. S7E). However, iDMG cells were routinely grown in the presence of EGF, which could potentially mask subtle, AREG-dependent influences on EGFR signaling. To dissect these differences, we cultured TH, TP, and THP iDMG with and without EGF and FGF in their normal growth media. All genotypes grew more slowly without growth factors (Fig. S7F), but THP and TH had smaller differences in doubling time between growth factor depleted and normal conditions than TP iDMG (Fig. 5F), suggesting that H3.3K27M is associated with an EGF-independent mechanism of proliferation. To determine whether this mechanism is EGFR-mediated, we inhibited EGFR signaling using neratinib in media with and without EGF. Removing EGF sensitizes THP but not TH or TP to neratinib, reducing the area under the neratinib 72-hour dose-response curve in only THP iDMG (Fig. 5G&H). This result suggests that THP iDMG are more dependent on EGFR pathway for maintaining their proliferative capacity than TH or TP iDMG. Taken together, these data suggest that the combination of H3.3K27M and PDGFRA^D842V^ induces an epigenetic program that upregulates amphiregulin expression and secretion, which correlates with growth factor-independent proliferation and enhanced sensitivity to EGFR tyrosine kinase inhibition.

### H3.3K27M and PDGFRA^D842V^ contribute to metabolic changes in tumor cells

Because THP iDMG upregulate proliferation-related genes and PDGFRA is upstream of several kinases, we investigated whether any druggable kinases are unique to THP. We first determined which kinases are significantly upregulated in iDMG vs. iNPC for each of the three genotypes. Seven kinases were upregulated only in THP tumors during the process of tumorigenesis (Fig. 6A). Of these, only *PCK2* was significantly overexpressed in THP relative to TH or TP iDMG, suggesting that it is important for both THP iDMG tumorigenesis and the unique biology of these tumors (Fig. 6B&C). Since *PCK2* has been shown to modulate anaerobic and aerobic metabolism in cancer (Vincent et al. 2015), we explored the impact of H3.3K27M and PDGFRA^D842V^ on cellular metabolism.

**Figure 6.**
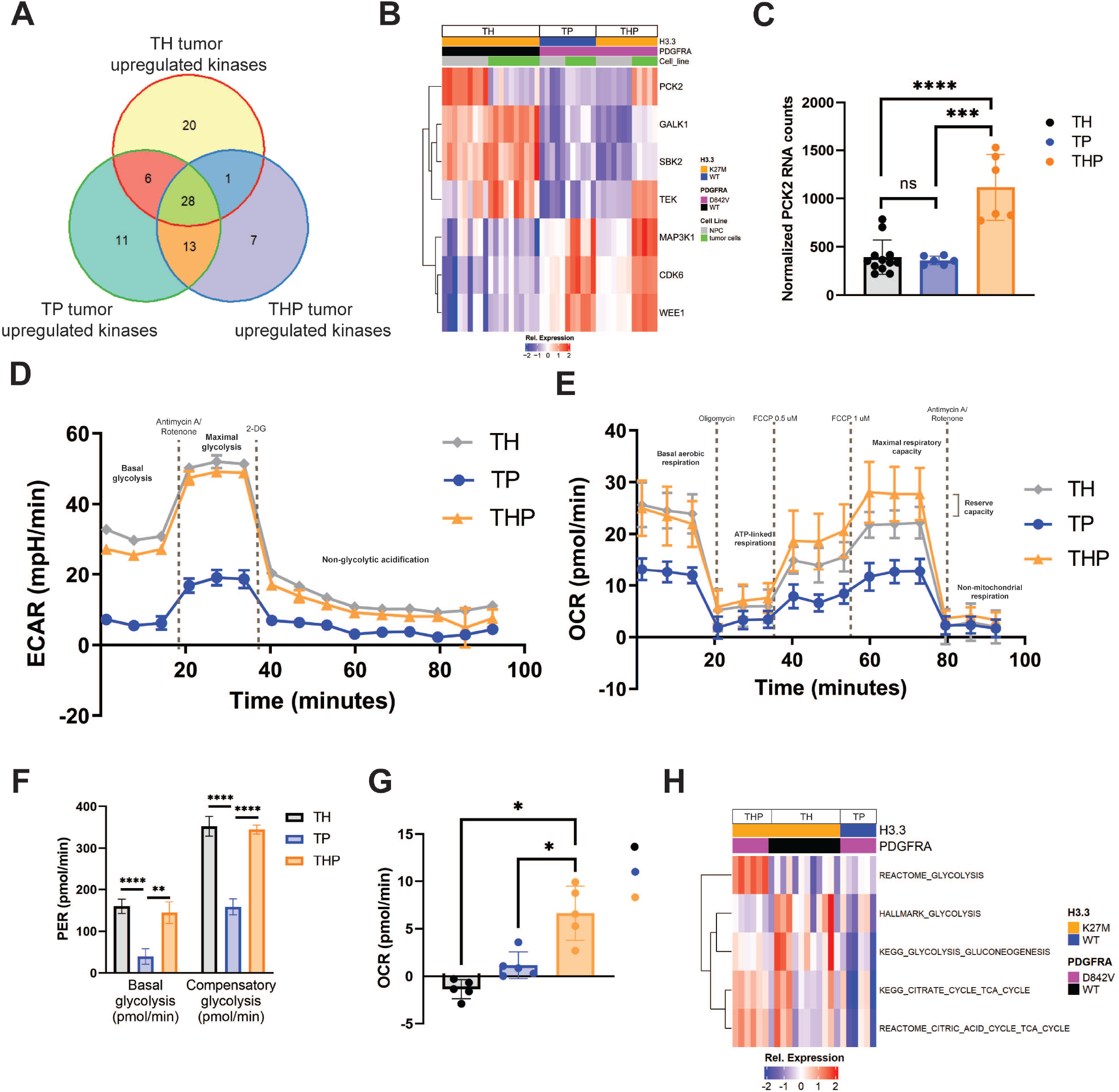
PDGFRA and H3.3 differentially affect both anaerobic and aerobic metabolism in iDMG. (**A**) Venn diagram of kinases significantly (Q<0.05) upregulated in TH, TP, and THP iDMG vs. their respective iNPC cells of origin. The 7 unique THP tumor upregulated kinases (purple section of the Venn diagram) represent targets that may be important in tumor initiation or maintenance only in THP iDMG. (**B**) Heatmap of iDMG normalized RNA expression of the 7 THP-specific kinases upregulated during tumorigenesis (from A). (**C**) DESeq2 normalized RNA counts of PCK2 (Welch’s t-test, ****=p<0.0001, ***=p<0.005). (**D**) Extracellular acidification rate of TH, TP, and THP iDMG cells in a Seahorse glycolytic rate assay showing an overall higher rate of glycolysis in H3.3K27M vs. WT iDMG. n=5 replicates per genotype. (**E**) Oxygen consumption rate of TH, TP, and THP iDMG cells in a Seahorse mitochondrial stress test showing differential rates of aerobic respiration between H3.3K27M and H3.3WT iDMG cells. n=5 replicates per genotype. (**F**) Glycolysis rates for TH, TP, and THP iDMG at baseline (basal) and maximum (compensatory) (t-test, **=p<0.01, ****=p<0.0001). (**G**) Reserve aerobic capacity calculated by subtracting baseline oxygen consumption from maximal oxygen consumption during the Seahorse mitochondrial stress test (t-test, *=p<0.01). n=5 replicates per genotype. (**H**) Relative enrichment scores for selected metabolism gene sets in iDMG cells. Scores were calculated using ssGSEA.

We profiled both glycolytic and aerobic metabolism of TH, TP, and THP iDMG cells *in vitro* using Seahorse assays. TH and THP had higher baseline rates of both glycolytic (Fig. 6D) and aerobic (Fig. 6E) metabolism than TP cells, consistent with previous studies showing that H3.3K27M increases general cellular metabolic activity (Chung et al. 2020). The H3.3K27M genotypes (TH and THP) also had significantly higher compensatory glycolytic rates than TP iDMG, indicating that they have a larger capacity to meet energetic demands (Fig. 6F). Both also had a higher maximal mitochondrial respiratory rate than their TP counterparts. However, THP cells were the only genotype exhibiting any reserve aerobic capacity, indicated by a maximal aerobic capacity greater than their baseline oxygen consumption (Fig. 6G). The robust aerobic and anaerobic metabolic capacities of THP cells suggest an ability to adapt to stressful environmental changes. Concurrently, THP iDMG express genes in both glycolysis and TCA cycle gene sets at a higher level than either TH or TP iDMG (Fig. 6H). These data suggest that the combination of PDGFRA^D842V^ and H3.3K27M alters the balance between aerobic and anaerobic metabolism and increases ability to adapt to energetic demand.

### H3F3A and PDGFRA status influence iDMG therapeutic responses

Given the reliance of THP iDMG on EGFR and PDGFRA signaling, we tested whether disruption of signaling downstream from these receptors using the blood-brain barrier penetrant MEK inhibitor trametinib would be effective. Trametinib is currently in a phase I/II clinical trial in pediatric glioma and has shown efficacy in H3.3K27M pHGG cell lines harboring MAPK pathway alterations (Izquierdo et al. 2022; Perreault et al. 2019). We found that both THP and TP, but not TH, iDMG are sensitive to trametinib, (Fig. S8A). This enhanced trametinib sensitivity is likely due to reliance on constitutive PDGFRA signaling through RAS/RAF/MEK for growth or survival. We also tested the sensitivity of iDMG to ONC201, a blood brain barrier-penetrant imipridone with anti-glioma activity *in vitro* and currently in a clinical trial in H3 K27-altered DMG (Allen et al. 2013; Arrillaga-Romany et al. 2017; Chi et al. 2019). All three iDMG were relatively ONC201-resistant, with more than 50% of cells surviving even at the highest drug doses (Fig. S8B). Given this resistance and the sensitivity of TP and THP iDMG to trametinib, we tested ONC201 and trametinib in combination. Trametinib and ONC201 synergistically inhibited growth of THP but not TH or TP iDMG cells, particularly at the highest ONC201 concentrations (Fig. 7A-C). These data suggest that this combination might prove effective in DMG with H3.3 and PDGFRA alterations.

**Figure 7.**
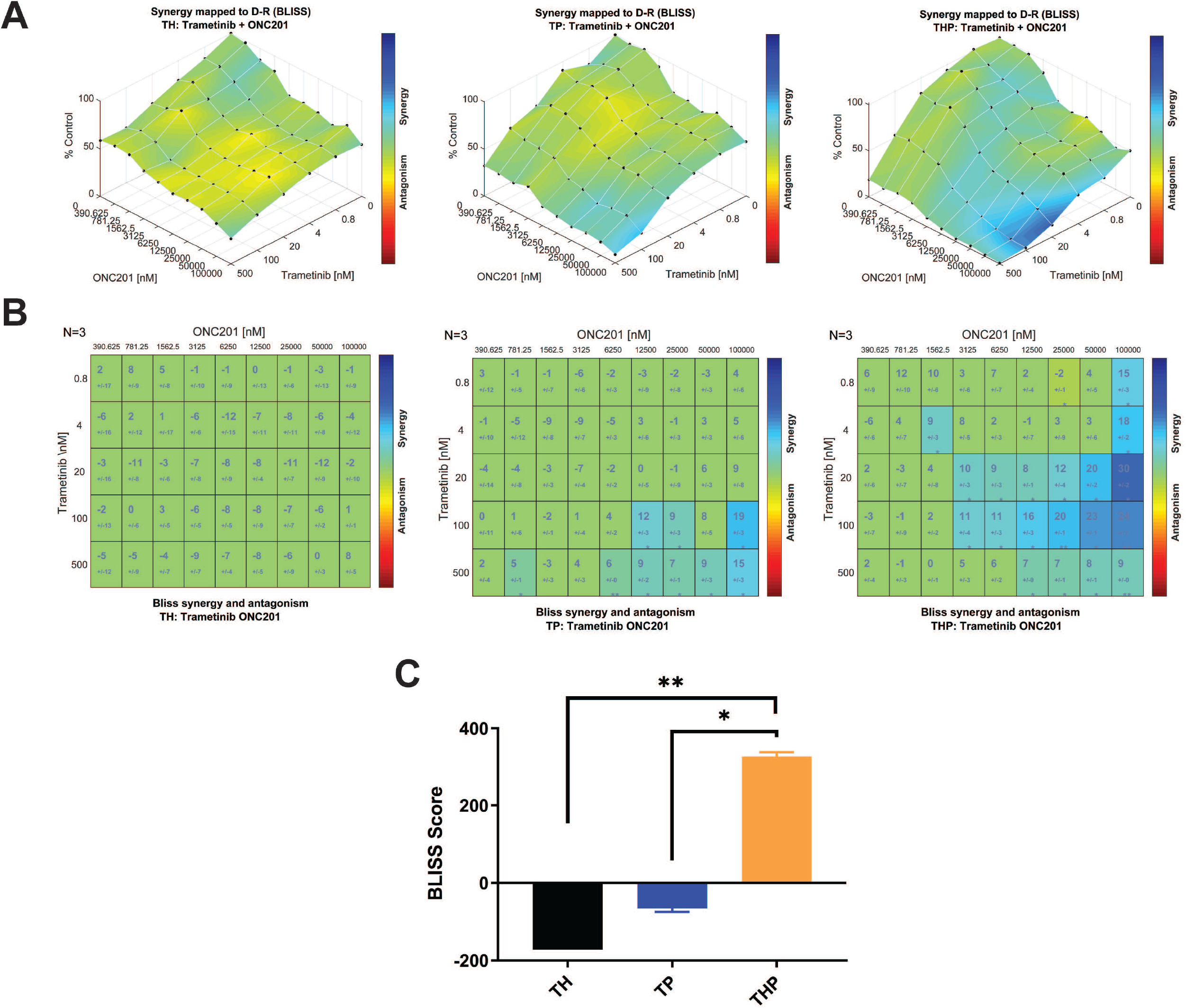
The combination of H3.3K27M and PDGFRA^D842V^ confers unique vulnerability to dual-agent therapy with ONC201 and trametinib. (**A**) Surface plots of BLISS synergy scores for dual ONC201/trametinib treatment in (left to right) TH, TP, and THP iDMG tumor cells. (**B**) BLISS synergy scores for each ONC201/trametinib combination in (left to right) TH, TP, and THP iDMG tumor cells. (**C**) BLISS synergy score sums (sum of all scores from the plots in B) for dual ONC201/trametinib treatment in each genotype. t-test, *=p<0.05, **=p<0.005.

## Discussion

*PDGFRA* alteration is a common event in H3.3K27M DMG, and its location upstream of the RAS/MAPK pathway makes it an attractive therapeutic target. However, its impact on the biology of these tumors is understudied. Here, we use a novel, iPSC-based model of H3.3K27M DMG to show that PDGFRA^D842V^ and H3.3K27M cooperate to increase tumor cell proliferation, alter intracellular signaling and metabolism, and present vulnerabilities to combination therapy. A variety of mouse and human model systems of H3.3K27M DMG have been used to elucidate the impact of this histone mutation on the epigenetic and transcriptomic landscapes of normal and tumor cells (Cordero et al. 2017; Grasso et al. 2015; Grigore et al. 2023; Haag et al. 2021; Larson et al. 2019; Misuraca et al. 2016; Pathania et al. 2017). iDMG augment these existing models in several important ways.

We demonstrated the potential of this model system by analyzing both iDMG and the iNPC from which they originated. IL6-JAK-STAT3 pathway gene upregulation is a characteristic of iDMG tumorigenesis regardless of *H3F3A* or *PDGFRA* genotype. These temporal analyses serve as a window into the changes that occur between normal brain cells and tumors. Such studies would be impossible in other human models such as PDX. Additionally, while our approach is similar to another iPSC-based model of H3.3K27M DMG (Haag et al. 2021), iDMG better recapitulate the genetics of patient tumors by harboring *PDGFRA* alteration alone and in combination with H3.3K27M. This approach showed that the specific combination of H3.3K27M and PDGFRA^D842V^ alters tumor biology in at least three important ways. First, it conferred an aggressively invasive, proliferative tumor phenotype by histology. While mice harboring THP iDMG had the same survival as those harboring TP iDMG, the THP tumors were larger and more invasive. The similar survival in mice with TP iDMG may be attributable to hydrocephalus caused by leptomeningeal tumor spread, which has been demonstrated in other mouse glioma models (Misuraca et al. 2016). Second, THP iDMG showed amphiregulin upregulation and consequent differential response to EGFR signaling manipulation. THP iDMG were resilient in terms of proliferation when deprived of exogenous growth factors, but were significantly sensitized to EGFR inhibition. Consequently, the combination of H3.3K27M and PDGFRA^D842V^ confers a unique reliance on the EGFR pathway that could be exploitable via strategies explored in adult GBM (Lin et al. 2022). Third, THP had a unique metabolic signature compared to TH or TP iDMG characterized by increased general metabolic rate, increased *PCK2* expression, and a slight elevation in reserve capacity. Recent work has shown that H3.3K27M expression increases both glycolytic and TCA cycle metabolism in neural progenitor cells (Chung et al. 2020), which is consistent with data showing increased basal glycolysis and oxygen consumption in TH and THP vs. TP iDMG. However, elevated reserve capacity and expression of *PCK2* are unique to THP iDMG and suggest an enhanced metabolic plasticity to respond to cellular energy demand and environmental stressors.

The genetic modularity of iDMG make them an ideal platform not only for isolating genotype-specific influences on tumor biology, but also for testing precision oncology-based therapeutic strategies. Studies dissecting the mechanism of action of ONC201 have shown that it may specifically target mitochondrial metabolism (Bonner et al. 2021), making it an interesting potential therapy to test in THP iDMG given their unique metabolic phenotype. However, the relative resistance of THP iDMG to single-agent ONC201 suggests that PDGFRA alterations may complicate clinical response. Previous work showed that gliomas with MAPK pathway alterations are susceptible to MEK inhibition (Izquierdo et al. 2022). Here, we found the combination of ONC201 and trametinib synergistically inhibits tumor cell growth only in THP iDMG. This finding raises two important points. First, better molecular stratification of patients for driver oncogene alterations will likely improve the efficacy of new therapies used in H3.3K27M DMG clinical trials. Second, dual-agent therapy based on driver oncogene profiles could be beneficial over single-agent therapies in H3K27-altered DMG patients (Louis et al. 2021).

Overall, this study validates the utility of iDMG as a model of patient tumors and interrogates the cooperativity between *PDGFRA* alteration and H3.3K27M. Our work demonstrates that H3.3K27M; PDGFRA altered tumors pose unique challenges, but also unique vulnerabilities. Moreover, it establishes a framework for using iDMG in mechanistic and therapeutic investigations that will hopefully improve the treatment of DMG patients.

## Materials and methods

### Cell culture

iPSC, iNPC, and iDMG tumor cells were cultured as previously described (Koga et al. 2020; Miki et al. 2022; Parisian et al. 2020). See Supplemental Methods for more details.

### Xenografting edited neural progenitors in mice

Animal experiments were approved by and performed under the regulations of the UCSD Animal Care Program (S00192M) and the University of Minnesota Institutional Animal Care and Use Committee (2003-37990A). iNPC were dissociated using Accutase (Innovative Cell Technologies), washed with PBS, and resuspended at 1 × 10^6^ cells in 2 μL PBS supplemented with 0.1% BSA per animal. Resuspended cells were kept on ice and inoculated into the pons of 4–6 week-old female Nod scid mice (Charles River Laboratory) by stereotactic injections at 1.5 mm to the right of midline, posterior to the lambdoid suture, and 5 mm deep from the inner skull plate (Aoki et al. 2012).

### Hematoxylin and eosin (H&E) staining and immunohistochemistry

Paraffin-embedded tissue blocks were sectioned and stained with H&E at the UCSD Moores Cancer Center Pathology Core.

### Cell growth and drug inhibition assays

Cell line growth and drug response were measured using either live-cell counting or CellTiter-Glo. See Supplemental Methods for details.

### Metabolism experiments

XF Cell Mito and XF Glycolysis Stress Test Kits were used to measure mitochondrial and glycolytic function on a Seahorse Extracellular Flux Analyzer (Agilent). The Seahorse microplate was coated with Cultrex (R&D Systems) for one hour at room temperature prior to cell plating. 10,000 cells per well were plated with 5 technical replicates per genotype. Assay medium was DMEM pH 7.4 with 2.5 mM glutamine, 17.5 mM glucose, and 1 mM pyruvate to mimic the composition of the normal tumor cell growth medium (DMEM/F12).

### Drug synergy assays

Two thousand cells of each cell line were plated in each well of Corning 96 well black microplates (Millipore Sigma) and treated with trametinib (Selleckchem) and/or ONC201 (Selleckchem) for 72 hours. ATPlite 1step (PerkinElmer) was used to measure the surviving fractions after the treatment in triplicate per manufacturer’s instructions. Synergy was evaluated using Combenefit (Di Veroli et al. 2016).

### RNA-seq library preparation

Total RNA was extracted from biological triplicate cell pellets using RNeasy Mini Kit (Qiagen) per manufacturer’s instructions. RNA libraries were constructed using KAPA mRNA HyperPrep Kit (KAPA Biosystems) and indexed with TruSeq single-index adapters to allow for sample multiplexing. Library quantity and quality were assessed using Qubit fluorometer (Thermo Fisher) and Agilent TapeStation 4150. Samples were randomly assigned to adapters and sequencing batches. Sequencing was performed using Illumina NextSeq 550 High Output Kits (v2.5, 75 cycles, single end).

### CUT&RUN library preparation

iDMG tumor cells were grown on Matrigel (Corning) in triplicate and harvested using Accutase. 250,000 cells per sample were frozen in individual aliquots in normal growth medium plus 10% DMSO. Prior to library preparation, frozen samples were quickly thawed in a 37° water bath and washed twice. Libraries were prepared using a CUTANA ChIC/CUT&RUN Kit (EpiCypher) per manufacturer’s instructions. Antibodies are listed in Supplemental Tables. E. coli DNA was spiked in at 0.5 ng/sample to permit quantitative comparison across samples. After antibody binding, Mnase digestion, and DNA purification, DNA libraries were constructed using the CUTANA CUT&RUN Library Prep Kit (EpiCypher) per manufacturer’s instructions. Library quantity and quality were assessed using a Qubit fluorometer (Thermo Fisher) and Agilent TapeStation 4150. All samples were pooled and sequenced using an Illumina NextSeq 550 High Output Kit (v2.5, 150 cycles, paired end).

### RNA-seq data analysis

FASTQ quality control was performed using fastQC and multiqc. RNA-seq reads were aligned to the human genome (v35) using STAR and reads were counted using Salmon (Dobin et al. 2013; Patro et al. 2017). To control for batch effects, selected samples were resequenced across experiments to ensure that they clustered according to biologic variables and not sequencing batch. Reads were filtered to remove low-count genes and transformed using variance-stabilized transformation in DESeq2. Differential expression analysis was performed using DESeq2 with a Benjamini-Hochberg-adjusted significance cutoff of Q<0.05 (Love et al. 2014). Single-sample gene set enrichment analysis was performed using GSVA. Gene set enrichment analysis was performed using GSEA and gene ontology analysis was performed using Gorilla and visualized using Revigo (Supek et al. 2011).

### CUT&RUN data analysis

Quality control was performed using fastQC and multiQC. Reads were aligned to both the *E. coli* and human genomes using bowtie2 (Langmead et al. 2012). Alignments were converted to paired-end BED files and then to normalized BEDGRAPHs (bedtools, samtools) using aligned *E. coli* reads as a scale factor (Li et al. 2009; Orlando et al. 2014; Quinlan et al. 2010). Peak calling was performed for each histone mark using SEACR in relaxed mode, comparing experimental samples to IgG negative controls (Meers et al. 2019). Peaks for each sample were merged for a given histone mark to make a consensus peak set. A table of read counts for consensus peaks was generated for downstream analysis (bedtools multicov). BigWig files were generated with deepTools bamCoverage and visualized using IGV (Ramirez et al. 2016). Heatmaps of consensus peak occupancy and average peak profiles were plotted using deepTools plotHeatmap. ChIP-Rx factors for H3K27me3 absolute amounts were calculated using the following formula:

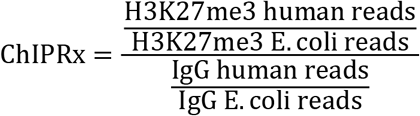

### Integration of gene expression and epigenetic analyses

RNA-seq and CUT&RUN data for a subset of iDMG samples were integrated using *intePareto* (Cao et al. 2020). This analysis calculates histone modification z-scores for each gene based on the relationship between the log fold change in RNA expression and histone mark occupancy, then maximizes z-scores for activating histone marks and minimizes them for repressive marks. H3K4me3 and H3K27ac were designated as activating histone marks and H3K27me3 was designated as repressive. *intePareto* plots were generated using ggplot2.

### Statistical analyses

All statistical analyses were performed using GraphPad Prism except for hypergeometric tests, which were performed on MSigDB. Data are representative of results obtained in at least 3 independent experiments.

## Data accessibility

Data have been deposited at with the following accession numbers: CUT&RUN (GSE225490) and RNA Seq (GSE225491) in Gene Expression Omnibus; patient-derived pHGG panel (EGAS00001002314, EGAS00001002328, EGAS00001004495, EGAS00001004496) in European Genome-Phenome Archive.

## Acknowledgments

This work was supported by R01NS080939 and R01NS116802 to F.B.F., the Japan Society for the Promotion of Science Overseas Research Fellowship to S.M., F31CA247177 and T32NS007431 to K.R.S., R01CA258248 to C.R.M and F.B.F., and R01CA204136 to C.R.M., T32GM008361 to B.L., and UL1TR001111 to C.R.M. and K.R.S. We thank Brittany Curtiss for advice on CUT&RUN analysis and Sriram Venneti for immunohistochemistry assistance. We are grateful to Kelley E. Smith-Johnston and Melissa J. Sammy for assistance with Seahorse experiments at the UAB Bioanalytical Redox Biology (BARB) Core, funded by P30DK079626, P30DK056336, UL1TR003096), CFRB and UCEM.

## Author contributions

T.K. and F.B.F. conceived this project and developed ideas in consultation with C.R.M. and R.S.H. T.K., S.M., K.R.S., F.N.G., and Y.S. performed wet laboratory experiments with input from E.T., D.M.S., D.M.M., R.J.W., E.N.B., B.C.M, and R.D.P. K.R.S, R.F.G., C.J., B.L., and A.M. performed computational analyses. E.H.T, D.M.S., and D.M.M. provided pathology services. T.K., S.M., K.R.S., and R.F.G. prepared the manuscript with supervision of C.C.C., R.S.H., C.R.M., and F.B.F. C.R.M. and F.B.F. secured funding to support this project and provided intellectual support for all aspects of the work. All authors contributed to manuscript revision.

