## Supplemental Methods for "Cooperativity between H3.3K27M and PDGFRA poses multiple therapeutic vulnerabilities in human iPSC-derived diffuse midline glioma avatars"

*Cell culture*

Experiments using human pluripotent stem cells were conducted under the regulations of the UCSD Human Research Protections Program, project number 151330ZX. Human induced pluripotent stem cell (iPSC) line iPS12-10 was purchased from Cell Applications. This cell line is integration-free and was validated for pluripotency, viability, karyotype normality, and normal disease status by Cell Applications. The iPS12-10 cells were cultured on plates coated with Matrigel hESC-Qualified Matrix (Corning) in mTeSR1 media (Stemcell Technologies). NPCs were cultured on Matrigel-coated plates in NPC maintenance media containing DMEM/F12 with GlutaMAX (Thermo Fisher Scientific), 1×N-2 supplement (Thermo Fisher Scientific), 1 ×B-27 supplement (Thermo Fisher Scientific), 50 mM ascorbic acid (Tocris), 3 μM CHIR99021 (Tocris) and 0.5μM purmorphamine (Tocris). Tumor cells were cultured on Matrigel (Corning) or Cultrex (R&D Biosciences) in DMEM/F12 with GlutaMAX (Thermo Fisher Scientific) with 1×B-27 supplement (Gibco), 20ng/ml EGF (Stemcell Technologies), and 20 ng/ml bFGF (Stemcell Technologies).

*Generation of genetically engineered human iPSCs*

pSpCas9(BB)-2A-GFP (px458) plasmid was a gift from Feng Zhang (Addgene plasmid #48138; http://n2t.net/addgene:48138; RRID:Addgene_48138) (Ran et al. 2013). The designated sgRNA sequences for each of the targeted genes were cloned into px458 using combinations of top and bottom oligonucleotides listed below.

TP53-R248Q-top: 5′ - CACCGCATGGGCGGCATGAACCGG - 3′

TP53-R248Q -bottom: 5′ - AAACCCGGTTCATGCCGCCCATGC - 3′

H3F3A-K27M -top: 5′ - CACCGAGAGGGCGCACTCTTGCGAG - 3′

H3F3A-K27M -bottom: 5′ - AAACCTCGCAAGAGTGCGCCCTCTC - 3′

Each pair of top and bottom oligonucleotides were phosphorylated and annealed by incubating 10 μM of each with 1 × T4 DNA ligase buffer (New England Biolabs), 5U T4 polynucleotide kinase (New England Biolabs) at 37 °C for 30 min, 95 °C for 5 min and by cooling down to 25 °C at 0.1 °C/s using a thermocycler. Annealed oligonucleotides were cloned into px458 by incubating 25 ng px458, 1 μM annealed oligonucleotides, 1× CutSmart buffer (New England Biolabs), 1 mM ATP (New England Biolabs), 10U BBSI-HF (New England Biolabs) and 200U T4 ligase (New England Biolabs) at 37 °C for 5 minutes, 23 °C for 5 min for 30 cycles. Correct cloning of each sgRNA sequence was confirmed by Sanger sequencing using U6 sequencing primer: 5′-GATACAAGGCTGTTAGAGAGATAATT-3′.

Single-stranded oligo DNA nucleotides (ssODNs) listed below were used to introduce the point mutation for each of the targeted genes.

TP53-R248Q-ssODN

5’-

TGACTGTACCACCATCCACTACAACTACATGTGTAACAGTTCCTGCATGGGCGGCAT

GAATCAACGCCCCATCCTCACCATCATCACACTGGAAGACTCCAGGTCAGGAGCCA

CTTGCCACCCTGCA - 3’

H3F3A-K27M-ssODN

5’-

TTTTCCTGTTTTTAATACCTGTAACGATGAGGTTTCTTCACCCCTCCAGTAGAGGGC

GCTGACATGCGAGCCGCTTTTGTAGCCAGTTGCTTCCTGGGTGCTTTACCACCGGTC

GATTTGCGGGCAGTCTGC - 3’

Human iPSCs were cultured in 10 µM Y-27632 RHO/ROCK pathway inhibitor for 2 h before dissociation. The cells were dissociated to single cells using Accutase (Innovative Cell Technologies). The dissociated iPSCs ($1 \times 10^6$ cells) were resuspended in 100 µl of supplemented solution of the Human Stem Cell Nucleofector Kit 1 (Lonza) containing a combination of the px458 plasmid targeting each gene and the ssODN and then electroporated using B-016 program of Nucleofector 2b (Lonza). The electroporated iPSCs were cultured on Matrigel-coated plates in mTeSR1 for 48 h. GFP-positive cells were then sorted by flow cytometer (SH800, SONY) and 1–$2 \times 10^4$ sorted cells were plated on a 10-cm Matrigel-coated plate in mTeSR1. Isolated colonies were manually picked and plated in duplicate Matrigel-coated 96-well plates.

The iPSCs clones on one of the duplicated 96-well plates were lysed using QuickExtract DNA Extraction Solution (Epicenter) and the PCR amplicons generated with the following primers were sequenced to confirm the edited iPSC clones.

TP53-intron6-forward: 5’ - GGGCCTGTGTTATCTCCTAG – 3’

TP53-intron7-reverse: 5' – GAGAGGTGGATGGGTAGTAG – 3'

H3F3A-intron1-forward: 5' – GCTGGTAGGTAAGTAAGGAG – 3'

H3F3A-intron2-reverse: 5' – GTTTTCCTGTTATCCATCTTTTTG – 3

*Differentiation of edited human iPSCs to neural progenitors*

Small molecule neural progenitor cells (smNPCs) were generated from iPSCs using an adapted protocol from a previous study (Reinhardt et al. 2013). In detail, human iPSCs at 70–80% confluency were dissociated using Accutase (Innovative Cell Technologies) and resuspended at $1\times10^6$ cells/ml in N2B27 medium (DMEM/F12 with GlutaMAX (Thermo Fisher Scientific), 1×N-2 supplement (Thermo Fisher Scientific), 1×B-27 supplement (Thermo Fisher Scientific), 150 mM ascorbic acid (Tocris), and 1% Penicillin/Streptomycin) supplemented with 1 μM Dorsomorphin (Tocris), 10μM SB431542 (Tocris), 3 μM CHIR99021, 0.5 μM Purmorphamine and 5 mM Y-26732 (Stemcell Technologies). Three million cells were transferred into one well of an uncoated six-well tissue culture plate and incubated at 37 °C, 5% $CO_2$ on a shaker at 90 rpm. Uniform small embryoid bodies (EBs) formed within 24 h and increased in size over the following days. After 48 h, a full media change was performed with N2B27 medium supplemented with Dorsomorphin, SB431542, CHIR99021, and Purmorphamine. At this time, about 2/3 of EBs were either discarded or split across three wells of a six-well plate to reduce the high cell density required initially to ensure uniform formation of EBs. On days 3–5, half media change was performed with fresh N2B27 media supplemented with Dorsomorphin, SB431542, CHIR99021, and Purmorphamine. On day 6, Dorsomorphin and SB431542 were withdrawn and a full media change with smNPC media (N2B27 media supplemented with 3μM CHIR99021 and 0.5μM Purmorphamine) was performed. At this stage, neuroepithelial folds were clearly visible in all EBs. On day 8, EBs were triturated by pipetting 10–15 times with a P1000 pipette and plated onto Matrigel-coated 10 cm plates. After 3 – 4 days, attached EB fragments and outgrown cells were dissociated to single cells with Accutase (Innovative Cell Technologies) and split at a 1:6 – 1:8 ratio onto Matrigel-coated plates. After the first passage, cells were passaged at a 1:10 – 1:15 ratio every 3 – 6 days.

*Lentivirus production and infection*

Human wildtype PDGFRA cDNA was subcloned into the pLV-EF1a-IRES-Puro lentiviral plasmid, a gift from Tobias Meyer (Addgene plasmid #85132; http://n2t.net/addgene:85132; RRID:Addgene_85132) (Hayer et al. 2016). The D842V mutation was introduced with site-directed mutagenesis using In-Fusion snap assembly master mix (Takara) according to the manufacturer's instruction. The primers listed below were used for the mutagenesis.

PDGFRA-D842V-forward: 5' - GGCCAGAGTCATCATGCATGATTCGAACTATGTG - 3'

PDGFRA-D842V-reverse: 5' - ATGATGCCTCTGGCCAGGCCAAAGTC - 3'

pLV-puromycin lentivirus expressing $PDGFRA^{D842V}$ was generated by co-transfection with VSVg and Δ8.9 packaging plasmids in 293T cells using Lipofectamine 2000 (Life Technologies). Supernatants were collected at 48 and 72 hours after transfection and virus was concentrated using Lent-X Concentrator (Takara). The NPCs were infected with lentivirus and subjected to 1 µg/mL puromycin selection.

*qRT-PCR*

RNA was extracted from cell pellets in triplicate using the RNeasy Mini Kit (Qiagen) according to the manufacturer's instructions and converted to cDNA using RNA to cDNA EcoDry premix (Takara). Reactions were run using the following primers:

GAPDH-RT-f: 5′-AATTTGGCTACAGCAACAGGGTGG-3′

GAPDH-RT-r: 5′-TTGATGGTACATGACAAGGTGCGG-3′

Nanog-RT-f: 5′-GAAATACCTCAGCCTCCAGC-3′

Nanog-RT-r: 5′-GCGTCACACCATTGCTATTC-3′

Oct4-RT-f: 5′-AGAACATGTGTAAGCTGCGG-3′

Oct4-RT-r: 5′-GTTGCCTCTCACTCGGTTC-3′

Pax6-RT-f: 5′-GCCCTCACAAACACCTACAG-3′

Pax6-RT-r: 5′-TCATAACTCCGCCCATTCAC-3′

Data were normalized to GAPDH expression.

*Cell viability assays*

For proliferation and drug toxicity experiments, cells were seeded on Cultrex (R&D Systems) in a 96-well plate at 500 cells/well. For proliferation experiments, viability was assayed using CellTiter-Glo 2D (Promega) every day for five days in technical quintuplicate. For drug toxicity, cells were treated with 9 drug concentrations plus a DMSO-only control in technical triplicates 24 hours after plating and assayed for viability using CellTiter-Glo 2D 72 hours after treatment. Briefly, CellTiter-Glo 2D reagent was added to assay wells at a 1:1 ratio with media and cell lysis was induced by agitation on an orbital shaker for two minutes. Luminescence was read ten minutes later using a Cytation 5 (Biotek). All cell viability assays were performed in biological triplicate.

*Live cell counting for proliferation*

iDMG were dissociated using Accutase and plated at various densities in technical sextuplicate on Matrigel-coated 96-well plates. Plates were incubated in a BioSpa 8 (Biotek) for 4.5 days and imaged on a Cytation 5 (Biotek) every 12 hours. Nine brightfield images of each well per timepoint were used for analysis. Image processing and cell counting scaled to the full area of the well were performed using Gen5 (Biotek, v. 3.10). Data analysis and curve fitting were performed using GraphPad Prism 9.1.2. All live cell imaging experiments were performed in biological triplicate.

*Western blotting*

Cells were dissociated using Accutase and lysed in RIPA buffer. Protein concentration was determined using a BCA kit according to the manufacturer's instructions (Pierce). Proteins (10 - 30 μg) were loaded on 4-15% gradient stain-free SDS-PAGE gels (BioRad). Blotting was performed using the TransBlot Turbo system (BioRad) and low-fluorescence PVDF membranes (BioRad) at 25V for 5-7 minutes. Membranes were blocked in 5% BSA for 1 hour at room temperature, then incubated with primary antibody at 4°C overnight (see Supplemental Tables for antibody information). Secondary antibody incubation (AlexaFluor 488 or 647, Thermo Fisher) was 30 minutes at room temperature. Membranes were imaged on the ChemiDoc MP (BioRad) and image analysis was performed using Image Lab (BioRad). Band intensities were normalized to total lane protein.
