## Supplemental Figures for "Cooperativity between H3.3K27M and PDGFRA poses multiple therapeutic vulnerabilities in human iPSC-derived diffuse midline glioma avatars"

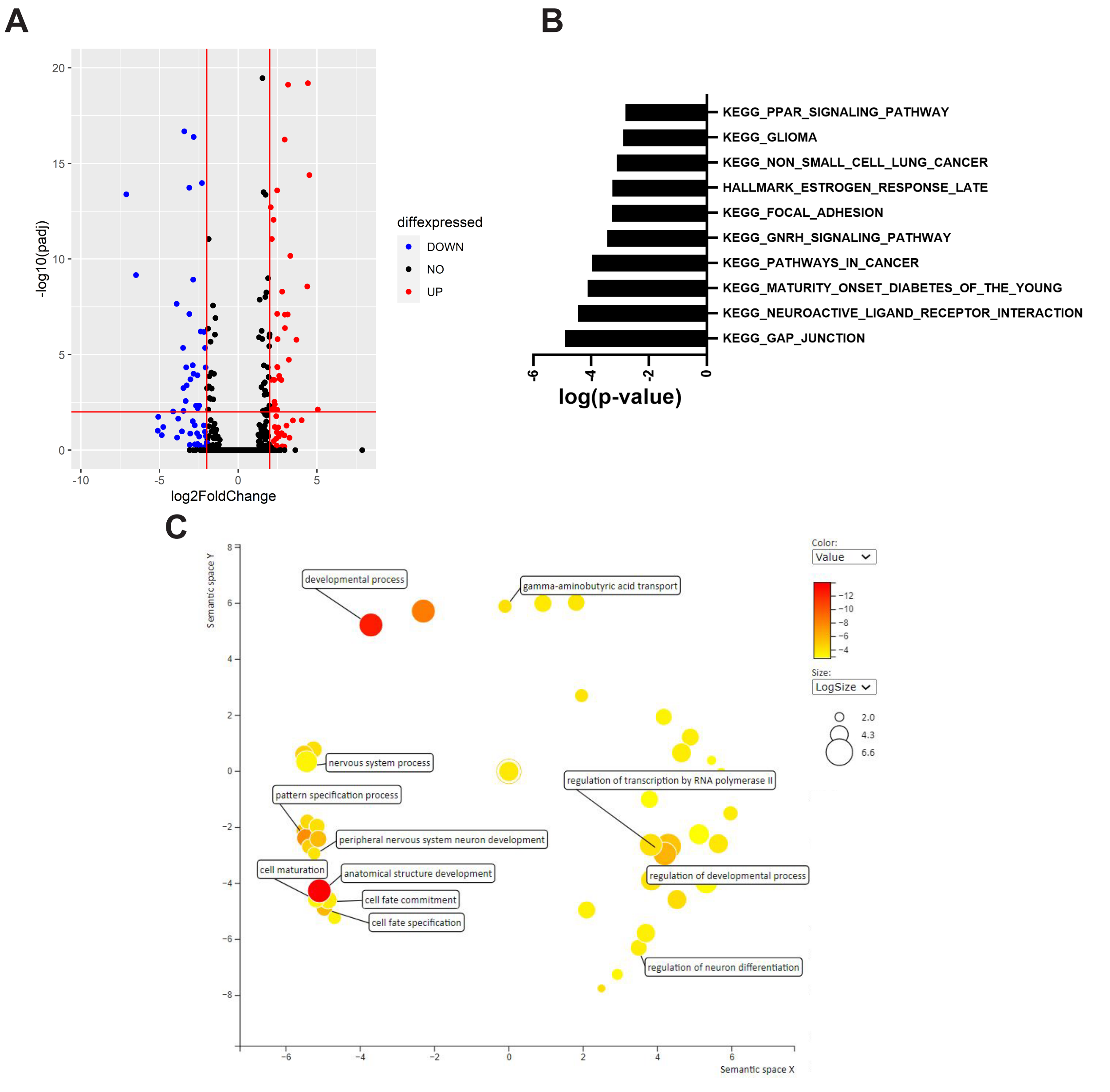
A
B
C
-log10(padj)
log2FoldChange
20
15
10
5
0
-10
-5
0
5
diffexpressed
DOWN
NO
UP
KEGG_PPAR_SIGNALING_PATHWAY
KEGG_GLIOMA
KEGG_NON_SMALL_CELL_LUNG_CANCER
HALLMARK_ESTROGEN_RESPONSE_LATE
KEGG_FOCAL_ADHESION
KEGG_GNRH_SIGNALING_PATHWAY
KEGG_PATHWAYS_IN_CANCER
KEGG_MATURITY_ONSET_DIABETES_OF_THE_YOUNG
KEGG_NEUROACTIVE_LIGAND_RECEPTOR_INTERACTION
KEGG_GAP_JUNCTION
-6
-4
-2
0
log(p-value)
Semantic space Y
Semantic space X
developmental process
gamma-aminobutyric acid transport
nervous system process
pattern specification process
regulation of transcription by RNA polymerase II
peripheral nervous system neuron development
cell maturation
anatomical structure development
regulation of developmental process
cell fate commitment
cell fate specification
regulation of neuron differentiation
Color:
Value
−12
−10
−8
−6
−4
Size:
LogSize
2.0
4.3
6.6

**Supplemental Figure S1. PDGFRA$^{D842V}$ enhances transcriptomic differences between H3.3K27M and H3.3WT iNPC.** (**A**) Volcano plot of gene expression changes between THP and TP iNPC. Red genes were significantly upregulated in THP cells, blue genes were significantly downregulated in THP cells, and black genes were not significantly differentially expressed. (**B**) Statistical significance of overlaps (hypergeometric test) between DEG in THP vs. TP iNPC and KEGG gene sets. (**C**) Gene ontology analysis of DEG between THP and TP iNPC visualized using ReviGO. Color represents log(p-value) and dot size represents the relative generality of the GO term.

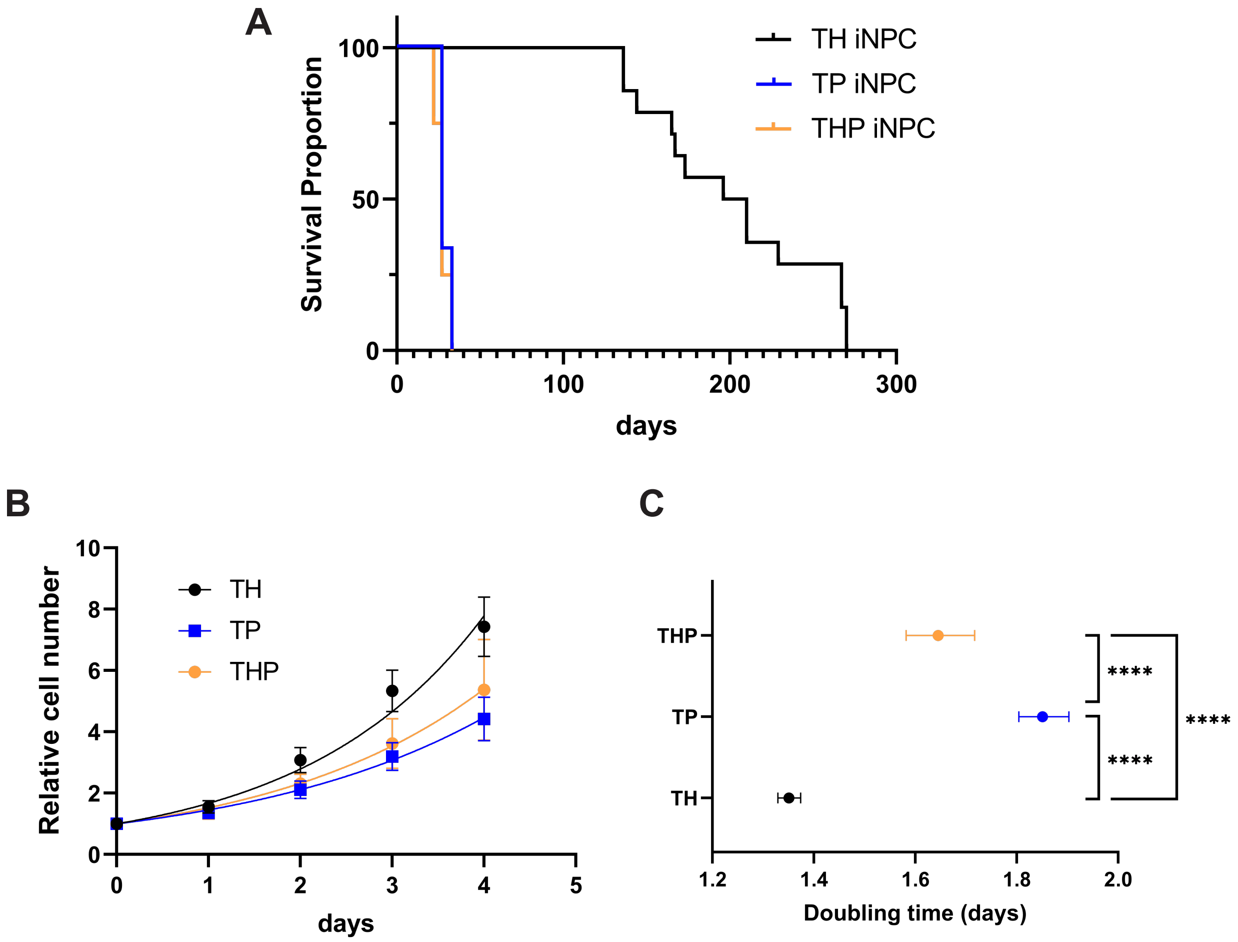
A
Survival Proportion
100
50
0
0
100
200
300
days
TH iNPC
TP iNPC
THP iNPC
B
Relative cell number
10
8
6
4
2
0
0
1
2
3
4
5
days
TH
TP
THP
C
THP
TP
TH
****
****
****
1.2
1.4
1.6
1.8
2.0
Doubling time (days)

**Supplemental Figure S2. iDMG grow robustly in tumor stem cell culture.** (**A**) Kaplan-Meier curves showing survival of immunodeficient mice orthotopically engrafted with TH, TP, or THP iNPC. Median survival of TH tumor bearing mice (n=14) was significantly longer than those bearing TP (n=3) or THP (n=4) tumors (log-rank test, $p<0.0001$). Median survival was not significantly different between mice bearing TP or THP tumors (log-rank test, $p=0.5936$). Survival curves were obtained from different experiments. (**B**) Proliferation of iDMG tumor cells *in vitro* measured using Cell Titer Glo. All measurements are normalized to day 0. n=2 experiments with 5 technical replicates each per genotype. (**C**) Doubling times of iDMG tumor cells *in vitro.* Error bars represent 95% confidence intervals calculated from the proliferation curves in A. Extra sum-of-squares F test, ****=$p<0.0001$.

A

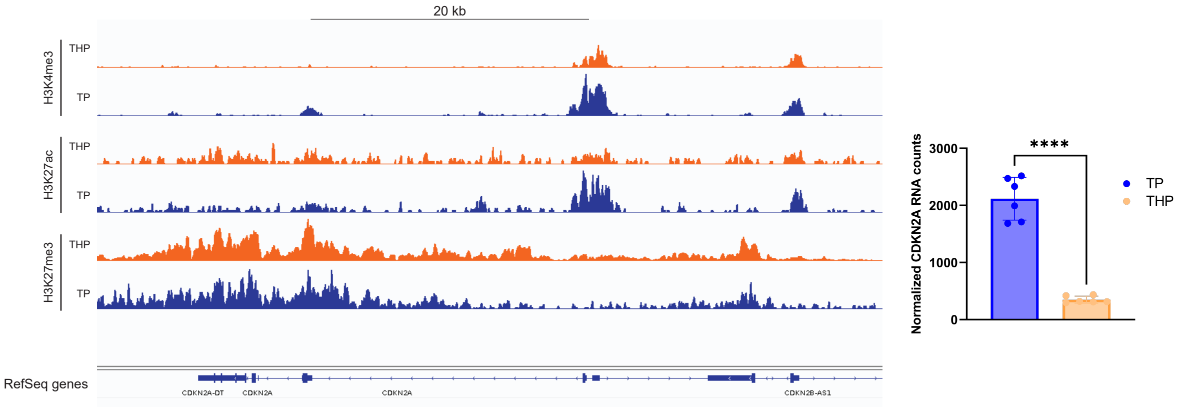

20 kb
H3K4me3
THP
TP
H3K27ac
THP
TP
H3K27me3
THP
TP
RefSeq genes
CDKN2A-DT
CDKN2A
CDKN2A
CDKN2B-AS1
Normalized CDKN2A RNA counts
3000
2000
1000
0
****
TP
THP


B

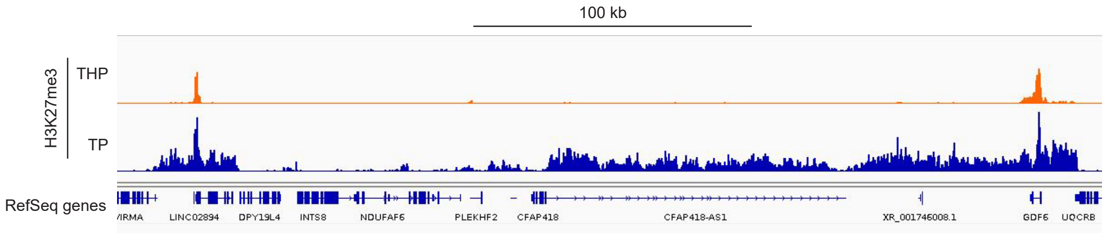

100 kb
H3K27me3
THP
TP
RefSeq genes
VIRMA
LINC02894
DPY19L4
INTS8
NDUFAF6
PLEKHF2
CFAP418
CFAP418-AS1
XR_001746008.1
GDF6
UQCRB

**Supplemental Figure S3. THP iDMG recapitulates epigenetic hallmarks of H3.3K27M expression.** (**A**) (Left) Representative H3K4me3, H3K27ac, and H3K27me3 cleavage under targets & release using nuclease (CUT&RUN) tracks for THP and TP iDMG cells at the *CDKN2A* locus show retention of H3K27me3 even with overall global H3K27me3 loss. This retention is accompanied by loss of H3K4me3 and H3K27ac. (Right) Normalized *CDKN2A* RNA counts show reduced *CDKN2A* expression in THP iDMG (t-test, ****=p<0.0001). (**B**) Representative CUT&RUN tracks for THP and TP iDMG cells showing reduced spreading of H3K27me3 from peaks.

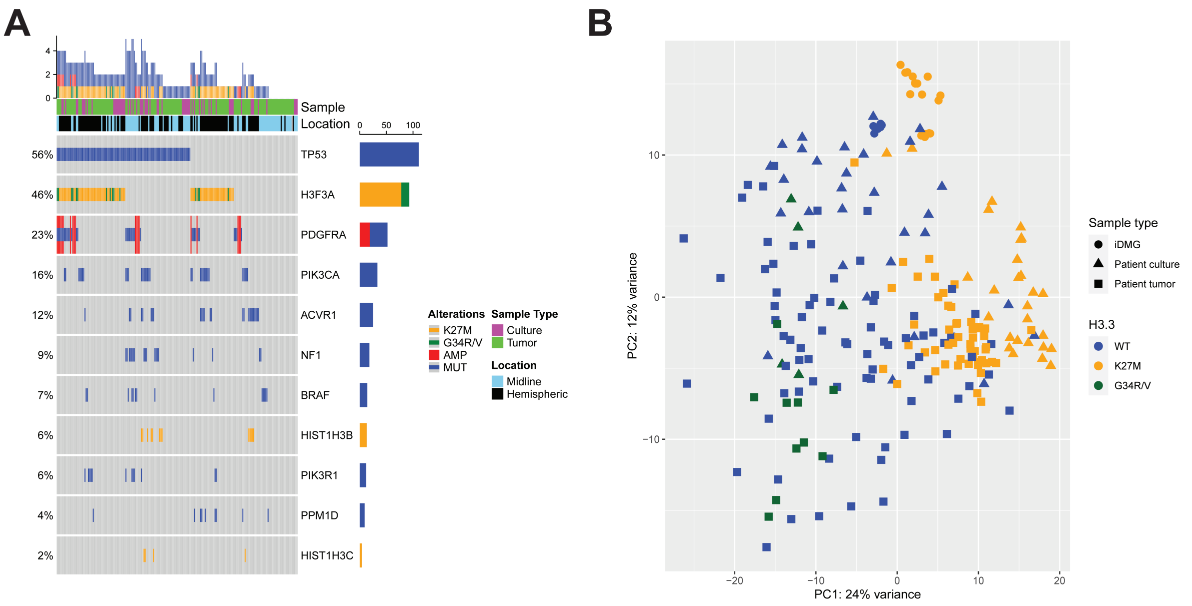


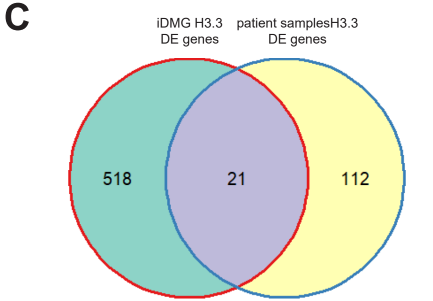


**D**

| Gene set | FDR q-value |
|---|---|
| GOBP_SENSORY_ORGAN_DEVELOPMENT | 3.71E-11 |
| GOBP_ANIMAL_ORGAN_MORPHOGENESIS | 3.22E-10 |
| GOBP_SENSORY_ORGAN_MORPHOGENESIS | 7.05E-09 |
| GOCC_CHROMATIN | 7.21E-08 |
| GOBP_SENSORY_SYSTEM_DEVELOPMENT | 8.87E-08 |
| DESCARTES_FETAL_CEREBRUM_INHIBITORY_NEURONS | 1.50E-07 |
| GOBP_TISSUE_DEVELOPMENT | 1.50E-07 |
| GOBP_EMBRYONIC_ORGAN_DEVELOPMENT | 1.50E-07 |
| BENPORATH_ES_WITH_H3K27ME3 | 2.10E-07 |
| GOBP_EMBRYO_DEVELOPMENT | 2.21E-07 |

**Supplemental Figure S4. THP iDMG are transcriptomically similar to H3.3K27M pHGG tumors and patient-derived cell lines.** (**A**) Oncoprint showing genetic alterations in a panel of pHGG tumors and patient-derived cell lines. The top bar graph shows total number of mutations in a given sample, while the right bar graph shows overall frequency of mutations in the sample panel. (**B**) Principal component analysis of RNA-seq data from iDMG and patient samples. Samples are clustered on 136 genes significantly differentially expressed ($P<0.05$) between H3.3K27M and non-K27M iDMG and patient samples. (**C**) Venn diagram of differentially expressed genes between H3.3K27M and non-K27M samples in iDMG (left) and patient samples (right) showing 21 genes commonly differentially expressed between the two sample groups (significant overlap via hypergeometric test, $p<6.581E-10$). (**D**) Top 10 most significant statistical overlaps between the 21 shared DE genes from (**C**) and all MSigDB gene sets.

A

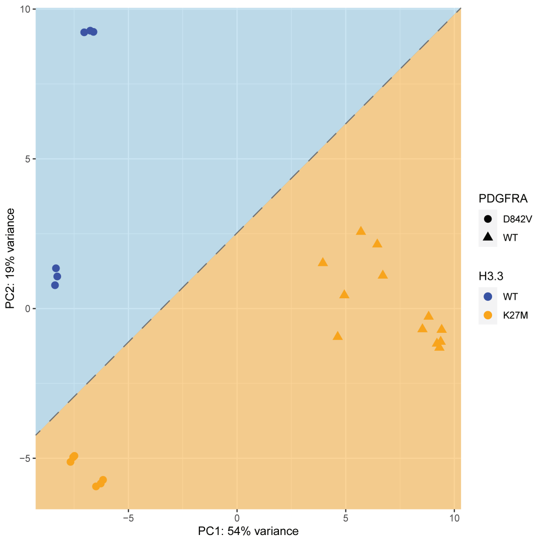


B

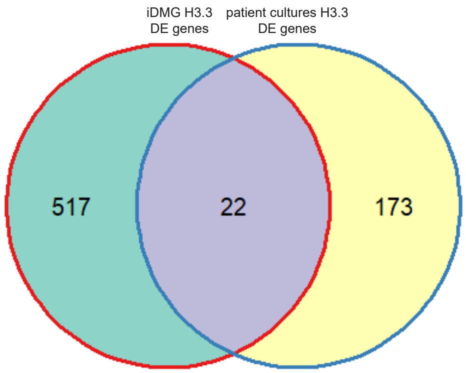


C

| Gene set | p-value |
|---|---|
| BENPORATH_ES_WITH_H3K27ME3 | 5.44E-11 |
| BENPORATH_EED_TARGETS | 2.92E-08 |
| GOBP_HEAD_DEVELOPMENT | 5.31E-08 |
| BENPORATH_PRC2_TARGETS | 2.05E-07 |
| BENPORATH_SUZ12_TARGETS | 3.27E-07 |
| GOBP_CENTRAL_NERVOUS_SYSTEM_DEVELOPMENT | 3.47E-07 |
| GOBP_SENSORY_ORGAN_DEVELOPMENT | 1.76E-06 |
| GOBP_REGIONALIZATION | 2.39E-06 |
| GOBP_NEGATIVE_REGULATION_OF_TRANSCRIPTION_BY_RNA_POLYMERASE_II | 2.73E-06 |
| GOBP_FOREBRAIN_DEVELOPMENT | 2.95E-06 |

**Supplemental Figure S5. iDMG share differential expression of PRC2 target genes with patient-derived cell lines based on H3.3 status.** (**A**) Principal component analysis of iDMG clustered on 190 genes significantly differentially expressed ($P<0.05$) between H3.3K27M and non-K27M pHGG patient-derived cell lines. (**B**) Venn diagram of differentially expressed genes between H3.3K27M and non-K27M samples in iDMG (left) and patient-derived cell lines (right) showing 22 genes commonly differentially expressed between the two sample groups (significant overlap via hypergeometric test, $p<9.279E-7$). (**C**) Top 10 most significant statistical overlaps between the 22 shared DE genes from (**B**) and all MSigDB gene sets.

**A**

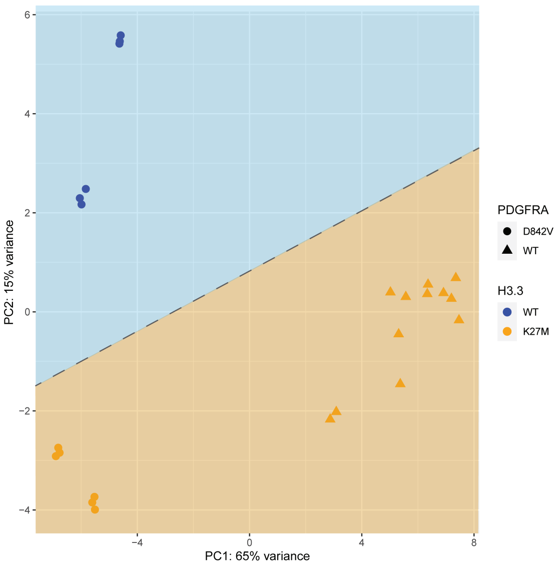


**B**

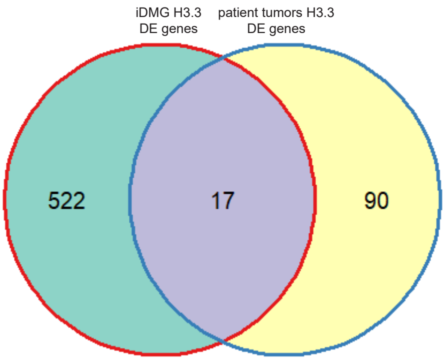


**C**

| Gene set | p-value |
|---|---|
| GOBP_SENSORY_ORGAN_DEVELOPMENT | 9.99E-07 |
| BENPORATH_ES_WITH_H3K27ME3 | 2.61E-06 |
| MIKKELSEN_MEF_HCP_WITH_H3K27ME3 | 2.35E-05 |
| DESCARTES_FETAL_CEREBRUM_INHIBITORY_NEURONS | 2.35E-05 |
| BENPORATH_EED_TARGETS | 2.35E-05 |
| MEISSNER_BRAIN_HCP_WITH_H3K4ME3_AND_H3K27ME3 | 2.35E-05 |
| FAN_EMBRYONIC_CTX_BIG_GROUPS_INHIBITORY | 6.16E-05 |
| ZHONG_PFC_MAJOR_TYPES_INTERNEURON | 1.99E-04 |
| GOBP_SENSORY_ORGAN_MORPHOGENESIS | 2.64E-04 |
| BENPORATH_SUZ12_TARGETS | 3.18E-04 |

**Supplemental Figure S6. iDMG share differential expression of PRC2 target genes with patient tumor samples based on H3.3 status.** (**A**) Principal component analysis of iDMG clustered on 107 genes significantly differentially expressed ($P<0.05$) between H3.3K27M and non-K27M pHGG tumors. (**B**) Venn diagram of differentially expressed genes between H3.3K27M and non-K27M samples in iDMG (left) and patient tumors (right) showing 17 genes commonly differentially expressed between the two sample groups (significant overlap via hypergeometric test, $p<1.253E-7$). (**C**) Top 10 most significant statistical overlaps between the 17 shared DE genes from (**B**) and all MSigDB gene sets.

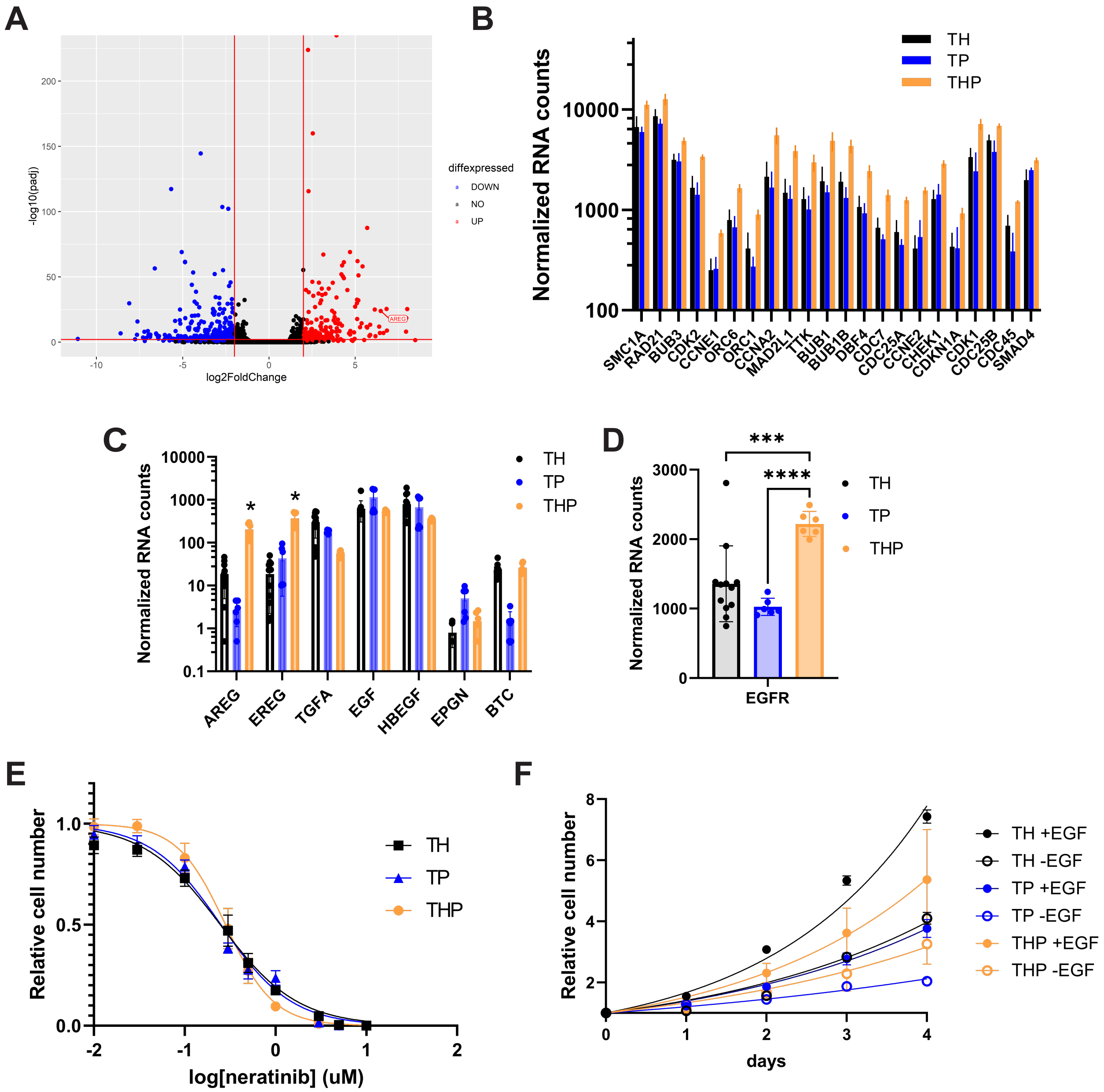

A
-log10(padj)
log2FoldChange
diffexpressed
DOWN
NO
UP
AREG
B
Normalized RNA counts
TH
TP
THP
SMC1A
RAD21
BUB3
CDK2
CCNE1
ORC6
ORC1
CCNA2
MAD2L1
TTK
BUB1
BUB1B
DBF4
CDC7
CDC25A
CCNE2
CHEK1
CDKN1A
CDK1
CDC25B
CDC45
SMAD4
C
Normalized RNA counts
AREG
EREG
TGFA
EGF
HBEGF
EPGN
BTC
D
Normalized RNA counts
EGFR
***
****
E
Relative cell number
log[neratinib] (uM)
F
Relative cell number
days
TH +EGF
TH -EGF
TP +EGF
TP -EGF
THP +EGF
THP -EGF

**Supplemental Figure S7. Increased AREG correlates with proliferation and sensitivity to EGFR pathway manipulation in THP iDMG.** A. Volcano plot of gene expression changes between THP and TP iDMG cells. *AREG* is significantly upregulated in THP vs. TP cells (log-2-fold change 6.47, Q=1.28e-24). B. Normalized RNA counts of genes in the KEGG Cell Cycle gene set that were uniquely upregulated in THP vs. TH and TP cells. Unique upregulation was determined by unsupervised hierarchical clustering of iDMG on the KEGG Cell Cycle gene set. All genes on the graph have significantly (t-test, $p<0.01$) higher expression in THP vs. TH and TP iDMG. C. Normalized RNA counts of EGFR ligands. *=THP counts are significantly greater than TH and TP counts, t-test, $p<0.001$. D. Normalized *EGFR* RNA counts. Welch's t-test, ***=$p<0.001$, ****=$p<0.0001$. E. Dose-response curve for neratinib toxicity in TH, TP, and THP iDMG. Relative cell numbers are normalized to zero drug, 1% DMSO control wells. n=9 replicates per genotype per concentration (3 experiments with 3 technical replicates each). F. *In vitro* proliferation curves for TH, TP, and THP iDMG cells in media with and without EGF and FGF. Relative cell numbers were normalized to day 0. n=6-12 replicates per genotype per condition (2-4 experiments with 3 technical replicates each).

A

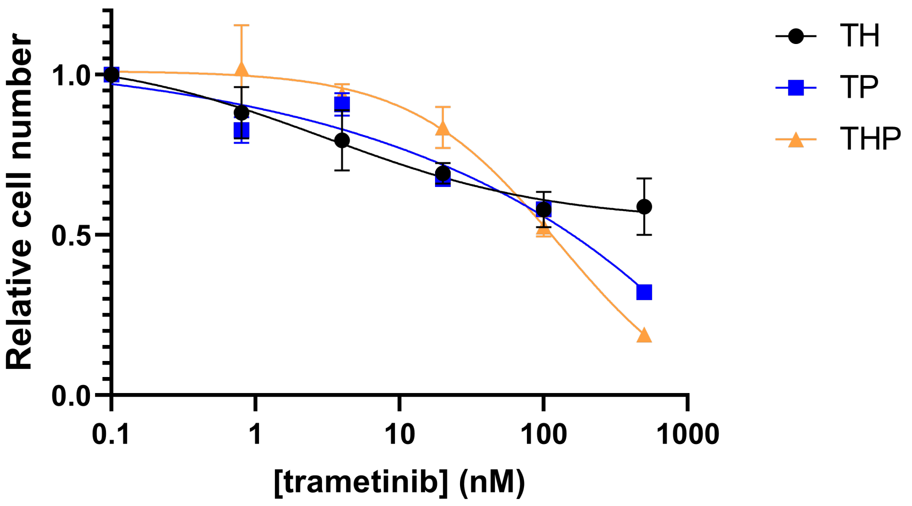

TH
TP
THP
1.0
0.5
0.0
Relative cell number
0.1
1
10
100
1000
[trametinib] (nM)


B

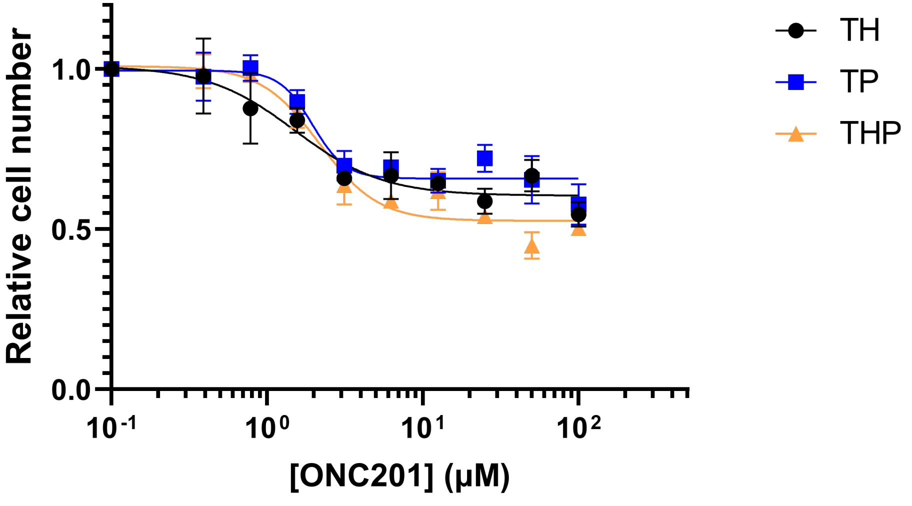

TH
TP
THP
1.0
0.5
0.0
Relative cell number
$10^{-1}$
$10^{0}$
$10^{1}$
$10^{2}$
[ONC201] (μM)

**Supplemental Figure S8. THP iDMG are sensitive to trametinib but resistant to ONC201.** Dose-response curves for trametinib (**A**) and ONC201 (**B**) toxicity in TH, TP, and THP iDMG. Relative cell numbers are normalized to zero drug, 1% DMSO control wells. n=3 biological replicates per genotype per concentration.
